# Collective Transitions from Orbiting to Matrix Invasion in 3D Multicellular Spheroids

**DOI:** 10.1101/2025.02.10.636936

**Authors:** Jiwon Kim, Hyuntae Jeong, Carles Falcó, Alex M. Hruska, W. Duncan Martinson, Alejandro Marzoratti, Mauricio Araiza, Haiqian Yang, Christian Franck, José A. Carrillo, Ming Guo, Ian Y. Wong

## Abstract

Coordinated cell rotation along a curved matrix interface can sculpt epithelial tissues into spherical morphologies. Subsequently, radially-oriented invasion of multicellular strands or branches can occur by local remodeling of the confining matrix. These symmetry-breaking transitions emerge from the dynamic reciprocity between cells and matrix, but remain poorly understood. Here, we show that epithelial cell spheroids collectively transition from circumferential orbiting to radial invasion via bi-directional interactions with the surrounding matrix curvature. Initially, spheroids exhibit an ellipsoidal shape but become rounded as orbiting occurs. However, cells gradually reorient from coordinated rotation towards outward strand invasion due to the accumulation of contractile tractions at discrete sites. Remarkably, the initial ellipsoid morphology predicts subsequent invasion of 2-4 strands roughly aligned with the major axis. We then perturb collective migration using osmotic pressure, showing that orbiting can be arrested and invasion can be reversed. We also investigate coordinated orbiting in “mosaic” spheroids, showing a small fraction of “leader” cells with weakened cell-cell adhesions can impede collective orbiting but still invade into the matrix. Finally, we establish a minimal self-propelled particle model to elucidate how collective orbiting is mediated by the crosstalk of cell-cell and cell-matrix adhesion along a curved boundary. Altogether, this work elucidates how tissue morphogenesis is governed by the interplay of collective behaviors and the local curvature of the cell-matrix, with relevance for embryonic development and tumor progression.

## 1. INTRODUCTION

Collective cell migration within a spherical geometry can occur along circumferential or radial directions, shaping tissue morphology via coordinated orbiting or matrix invasion [1]. For instance, normal epithelial cells undergo coherent rotational motions to organize spherical acini or alveoli when embedded within 3D matrix [2–8]. Moreover, follicle epithelial cells drive global egg chamber rotation within a basement membrane during *Drosophila* oogenesis [9, 10]. Analogous rotational motions have been observed when epithelial cells are confined in planar geometries within circular corrals [11–14] or migrating along cylindrical geometries [15]. In these scenarios, circumferential migration is guided by the curvature of the confining material, which remains largely static since cells cannot alter the local curvature. Thus, the effect of dynamic reciprocity, whereby cell migration is shaped by the matrix interface and the matrix interface is (in turn) shaped by cell migration, remains poorly understood.

Multicellular spheroids can also invade into the surrounding matrix by opening up cell-sized gaps using matrix metalloproteinases (MMPs) or irreversible deformation [16–25]. Such invasion may be facilitated by “leader cells” with enhanced capabilities for path generation or coordination of followers [18–20, 24–26]. Such leader cell phenotypes may be a consequence of the epithelial-mesenchymal transition (EMT), which is associated with a weakening of cell-cell adhesions, as well as front-back polarity with strengthened cell-matrix adhesions [27]. Indeed, in heterogeneous populations, cells with more mesenchymal states may sort away from cells with more epithelial states based on differences in cell-cell adhesion (e.g. E-cadherin). Historically, cell sorting was studied using embryos and spheroids in the absence of exogeneous matrix [28]. Nevertheless, cell sorting can be further mediated by differences in both cell-cell and cell-matrix adhesion when spheroids are embedded in 3D matrix [26, 29]. Computational models based on self-propelled particles with varying adhesion [30] can exhibit such self-sorting [31, 32], as well as coordinated motility (also described as flocking or milling) [33–35].

Here, we show that multicellular spheroids of mammary epithelial cells transition from collective orbiting to strand invasion based on dynamic mechanical interactions with the surrounding 3D matrix. Initially, multicellular spheroids are slightly ellipsoidal but become rounded due to coordinated rotational motion along the curved matrix interface. Subsequently, localized matrix deformations disrupt collective orbiting and interfacial curvature, promoting radial strand invasion at discrete locations along the periphery. Remarkably, these discrete sites of strand invasion are roughly aligned with the initial elongation axis of the spheroid. We reversibly manipulate these interfacial dynamics using osmotic pressure, which can drive retraction of strands back into the spheroid, as well as reversibly arrest orbiting. Further, we examine whether heterogeneous “mosaic” spheroids continue to orbit and invade when mammary epithelial cells are mixed with increasing fractions of EMT-induced mammary cells with weakened cell-cell adhesion. Finally, we develop an agent-based model that recapitulates collective transitions from orbiting to invasion based on geometric perturbations of local boundary curvature at a few discrete locations. Overall, this work reveals the dynamic interplay of collective migration and local curvature at a spheroid-matrix interface, with implications for epithelial tissue morphogenesis in development and disease.

## II. RESULTS

### Multicellular Spheroids Transition from Orbiting to Invasion in Collagen I Matrix

We investigated transitions between different collective cell migration modes using multicellular spheroids of mammary epithelial cells (MCF-10A) embedded within 3D collagen I hydrogels. Briefly, hanging drop culture was used to aggregate ∼ 500 cells into spheroids of ∼ 150 *µ*m in diameter, which were then embedded in 6 mg/mL bovine collagen I for live cell imaging (Fig. S1A-F), which exhibits a typical fiber spacing of 0.7 *µ*m (Fig. S1G-K). MCF-10A cells were stably transfected so that their nuclei were fluorescent (H2B-mCherry) and imaged using spinning disk confocal microscopy in an environmentally controlled chamber.

Cells within the spheroid exhibited collective orbiting motions within 6 h of embedding, continuing for the next 6 h (Fig. 1A, i-iv). Coordinated rotational motion primarily occurred in the focal plane based on 3D tracking of representative nuclei, so our analysis focused on cell motion in the equatorial plane of the spheroid (Fig. S2). We visualized rotational motion by unwrapping 20 *µ*m-thick concentric layers of cell (nuclei) at sequential time points as a kymograph (Fig. 1BC; S3AB). During the first 6 hours, the cell nuclei trajectory stayed parallel and horizontal, representing that the relative angular location of the cells are hardly changed (Fig. 1C, i-ii). Notably, the parallel upward trajectories from 6–12 h indicate highly coordinated angular motility within the outermost layer (Fig. 1C, ii-iv). Subsequently, cell nuclei exhibited flat trajectories through the next 24 h, indicating that their angular positions within the outermost layer were largely unchanged (Fig. 1C, iv-vi). Collective migration was then analyzed using optical flow, revealing sustained circumferential velocity fields in the outermost 40 *µ*m of the spheroid, which was less pronounced in the innermost 100 *µ*m of the core (Fig. 1D, ii-iv; S3C). In comparison, the radial components of the velocity were relatively uncorrelated and sporadic from 0-11 h (Fig. 1E, i-iv). However, cells located at the top and lower left of the spheroid veered radially after 12 h, disrupting circumferential migration (Fig. 1A, v). Indeed, “hotspots” of increased radial velocity were observed at 17 h, localized at the top and lower left (Fig. 1E, v; S3D). The spheroid morphology at the top and lower left became slightly sharper relative to the otherwise rounded periphery elsewhere. Subsequently, multicellular strand invasion was observed at the top and lower left by 24.75 h (Fig. 1A, vi), also associated with increased radial velocities (Fig. 1E, vi). These bursts of radially oriented velocity were very apparent in the outermost cells (Fig. S3F).

**FIG. 1.**
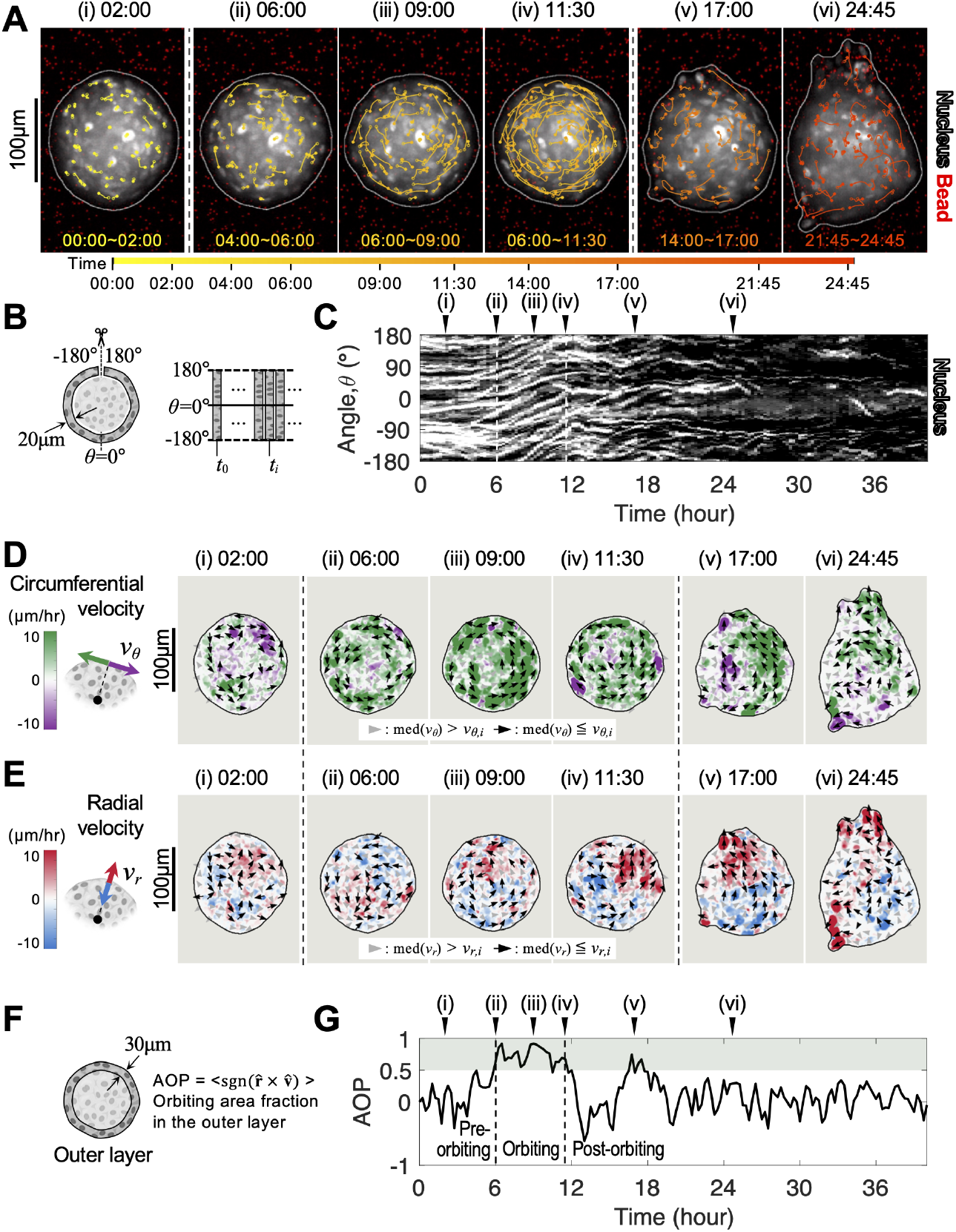
Multicellular spheroids transition from circumferential orbiting to radial invasion in 3D matrix. (**A**) Representative images of the MCF-10A spheroid embedded in a 3D collagen matrix, overlaid with selected trajectories of tracked nuclei. Trajectories are color coded from yellow to red to indicate different starting times over 24 h. Open and solid circles represent the starting and ending points, respectively. (**B**) Schematic of unwrapping and concatenating the outermost 20 *µ*m peripheral “layer” of the spheroid equatorial plane at each time to generate a kymograph. (**C**) Kymograph of fluorescent nuclei (H2B-mCherry) in the outermost peripheral layer of cells shows minimal angular motion from 0-6 h after embedding (i-ii). Next, all nuclei exhibit coordinated angular motion from 6-12 h, indicated by the parallel upward trajectories (iii-iv). Finally, nuclei exhibit minimal and uncoordinated angular motion as they transitioned into radial migration mode (v-vi). (**D**) Circumferential velocity profiles of the spheroid extracted with optical flow. The magnitude of the angular velocity is color coded from purple to green. Black arrows denote large velocities greater that the median, while gray arrowheads denote smaller velocities less than the median. (**E**) Radial velocity profiles of the spheroid extracted with optical flow. The magnitude of the angular velocity is color coded from blue to red. Black arrows again denote large velocities greater that the median, while gray arrowheads denote smaller velocities less than the median. (**F**) Angular order parameter (AOP) is based on the fraction of circumferentially migrating cells in the outermost 30 *µ*m peripheral “layer” of the spheroid equatorial plane at each time. (**G**) AOP values exceed 0.5 from 6-12 h during coordinated orbiting, but are below this value before and after.

The beginning and end of the orbiting phase were determined by 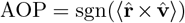, which describes the fraction of velocity pixels oriented circumferentially in the outermost region (Fig. 1F). We set a threshold of AOP greater than 0.5 for coordinated orbiting. From 6-12 h, AOP ≈ 0.75, indicating the high fraction of pixels within the designated peripheral area oriented to the primary orbiting direction (counterclockwise) (Fig. 1G, ii-iv). After 12 h, AOP decreased in magnitude with fluctuations comparable to the pre-orbiting period (Fig. 1G, iv-v). Although orbiting was most pronounced in the outermost layer, cells towards the interior also exhibited coordinated motion (Fig. S3B). Spheroid area remained roughly constant during orbiting but increased dramatically when invasion occurred (Fig. S3G).

### Curvature of Spheroid-Matrix Interface Shapes Collective Migration Mode

Next, we investigated how collective migration was shaped by the local curvature along the spheroid periphery, based on our observation that radial invasion proceeded from sharp outgrowths that perturbed the locally rounded morphology. Unexpectedly, we observed that most spheroids were ellipsoidal in the hanging drop and remained so after being embedded in collagen I (Fig. S1 B-F). As a representative example, a spheroid was slightly elongated (vertically) from 0-3 h, which we quantified based on the radial deviation from an equivalent reference circle, Δ*r* = 8 *µ*m at *θ* = 0^°^, oriented for convenience from the position of maximum protrusion at time = 0 h (Fig. 2AB, i). This spheroid was also slightly flattened (horizontally) with Δ*r* = -3 *µ*m on the left and right (*θ* = ±90^°^). The spheroid became more rounded as orbiting commenced, so that the radial deviation decreased to Δ*r* = 0 *µ*m at *θ* = 0^°^ and to Δ*r* = 1 *µ*m at *θ* = ±90^°^ from 8.5-10.5 h (Fig. 2AB, ii). We further quantified this based on roundness, 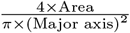, which is 1 for a perfect circle and approaches 0 for non-circular shapes. Typically, spheroids had a roundness of 0.9 prior to orbiting (0-6 h), which increased to almost 1.0 during orbiting (6-12 h) (Fig. 2C, i-ii). Typically, orbiting began at 5± 2 h and completed at 13 ± 2 h, lasting about 7.5 hours (Fig. 2D). However, the shape compaction, defined as the duration during which the roundness ranks in the top 10% within the first 24 hours, was observed slightly later, from 8 ± 3 h to 16± 3 h, lasting for 9 hours. Thus, coordinated orbiting typically preceded the morphological rounding of the spheroid.

**FIG. 2.**
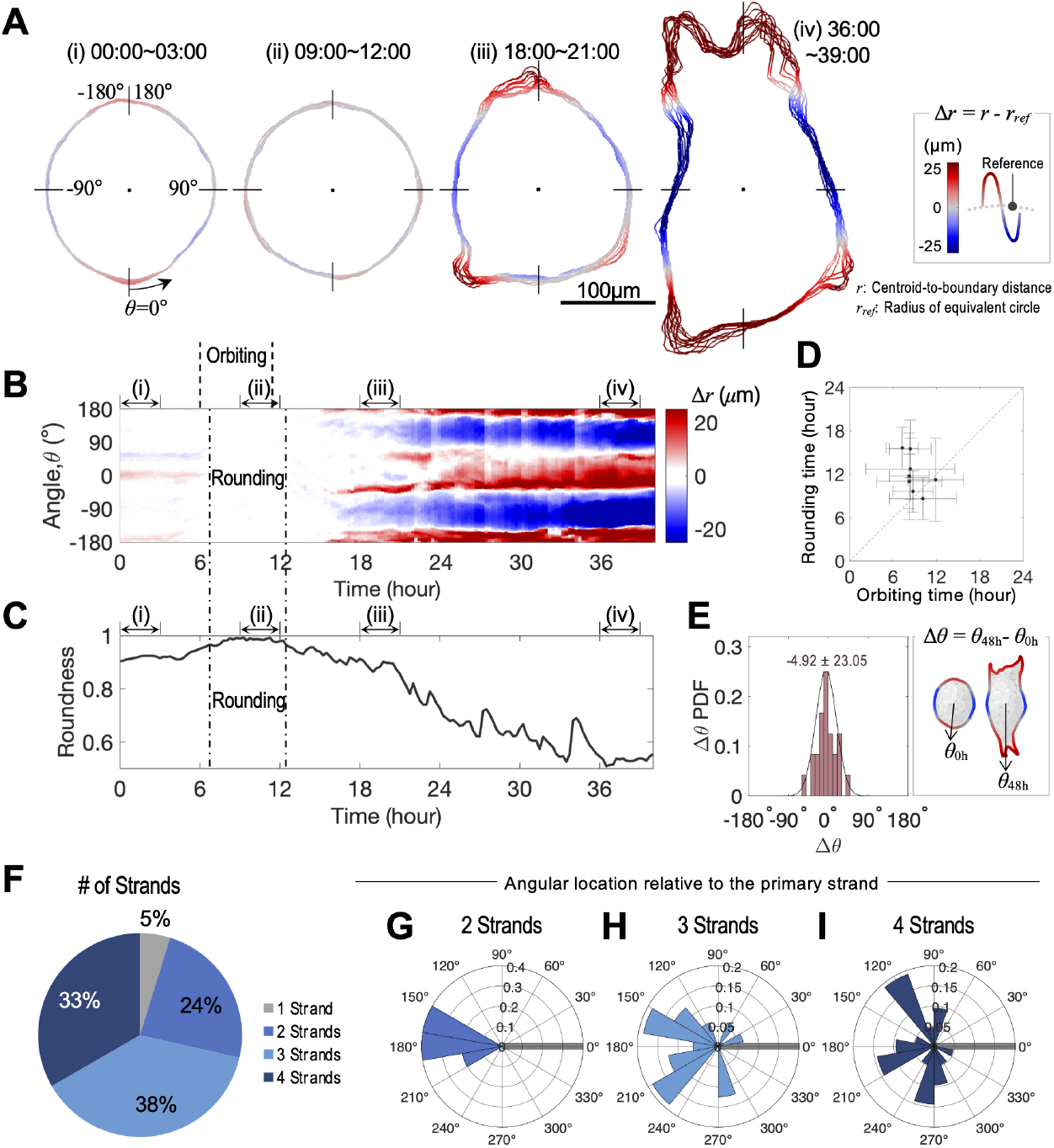
Spheroid morphology is initially elongated, rounding up during orbiting, then elongating again along the same axis for strand invasion. **(A)** Representative spheroid morphology based on cross-section at equatorial axis. The boundary is color-coded by larger (red) or smaller (blue) radial deviation from a reference radius for a perfect circle (gray). The references are set to the equivalent circle at each time point. During phase (ii) 09:00 ∼ 12:00, the boundary color is mostly gray, indicating that the spheroidal morphologies are comparable to the equivalent circles. **(B)** Kymograph of radial deviation along the periphery at each time, highlighting initial and final elongation sites. **(C)** Increase in roundness value from 6-12 h and decrease thereafter is comparable to orbiting and invasion. **(D)** Comparison of rounding time and orbiting time shows orbiting slightly precedes rounding. *n*_spheroids_ = 8 across duplication. **(E)** Comparison of angular positions of initial (00:00) and final (48:00) spheroid elongation axis. *n*_spheroids_ = 21 across triplication. **(F-I)** Number and angular distribution of multicellular strands. Solid gray lines at *θ* = 0^°^ indicate the primary (longest) strand. *n*_spheroids_ = 21 across triplication.

As the spheroid transitioned to matrix invasion after 12 h, there was a gradual increase in locally sharpened features at *θ* = 0^°^ and 180^°^ (Fig. 2AB, iii). This “roughening” of the periphery corresponded to a decrease of roundness to 0.8 (Fig. 2C, iii). Further, the invasion of multicellular strands from these angular locations corresponded to the increasing local curvature at *θ* = 0^°^ and 180^°^ at 36-39 h (Fig. 2AB, iv) and a further decrease in roughness to 0.5 (Fig. 2C, iv). Remarkably, the angular locations of invasive strands was highly correlated with the initially protruded poles of the spheroids (Fig. 2B, i, iii-iv). The angular positions of the most elongated region at 0 h and 48 h differed by as little as Δ*θ* = − 5 ± 23^°^ (Fig. 2E). Further, spheroids typically exhibited 2-4 invasive strands, aligned roughly with the initial elongation axis (Fig. 2F-I).

Since the multicellular spheroid was adherent to collagen I, it was hypothesized that alterations in spheroid morphology were driven by cell-generated forces that would further deform the surrounding 3D matrix. Optical flow analysis was used to track the displacement of far red fluorescent tracer microparticles embedded in the collagen network relative to their initial position (Fig. 3AB, S4). For example, the representative spheroid from Fig. 2 exhibited locally increased inward tractions at top and bottom relative to the rest of the periphery at 2 h (Fig. 3C, i), consistent with slight flattening at these locations (Fig. 2 A, i). During collective orbiting from 6-12 h, the spheroid exhibited a further increase in inward tractions at the top and bottom (Fig. 3C, ii-iv), even as the rounding of the boundary was nearly complete (Fig. 2A and C, ii). Moreover, there were subtle protrusive tractions at ± 90^°^, consistent with slight rounding (Fig. 3C, ii-iv). This orbiting regime was associated with a sizable increase in the amplitude of radial displacement undulations, indicating enhanced anisotropic deformation, (Fig. 3D). Moreover, confocal reflectance imaging indicated local collagen fiber alignment at these sites of large matrix displacement (Fig. 3E-H). To visualize these locally non-uniform matrix deformations, we modified our previous DART protocol to bin the displacement field at discrete angular positions [36]. Notably, this analysis indicates the increased matrix displacements at the *θ* = 0^°^ and 180^°^ of the spheroid from 6–9 h, which remain comparable until 12 h (Fig. 3I, ii-iii). Nevertheless, the transition to invasion after 12 h was associated with monotonically increasing contractile tractions near 0^°^ and 180^°^ (Fig. 3I, v-vi, S5). For comparison, bead displacement computed non-cumulatively at sequential 2 h intervals revealed large fluctuations in matrix displacement at varying angular positions along the periphery, mostly during orbiting (Fig. 3J-L, S6). Overall, these results show that transitions from circumferential orbiting to radial invasion are governed by the buildup of matrix deformation which then perturbs the local curvature of the spheroid periphery (Fig. S7).

**FIG. 3.**
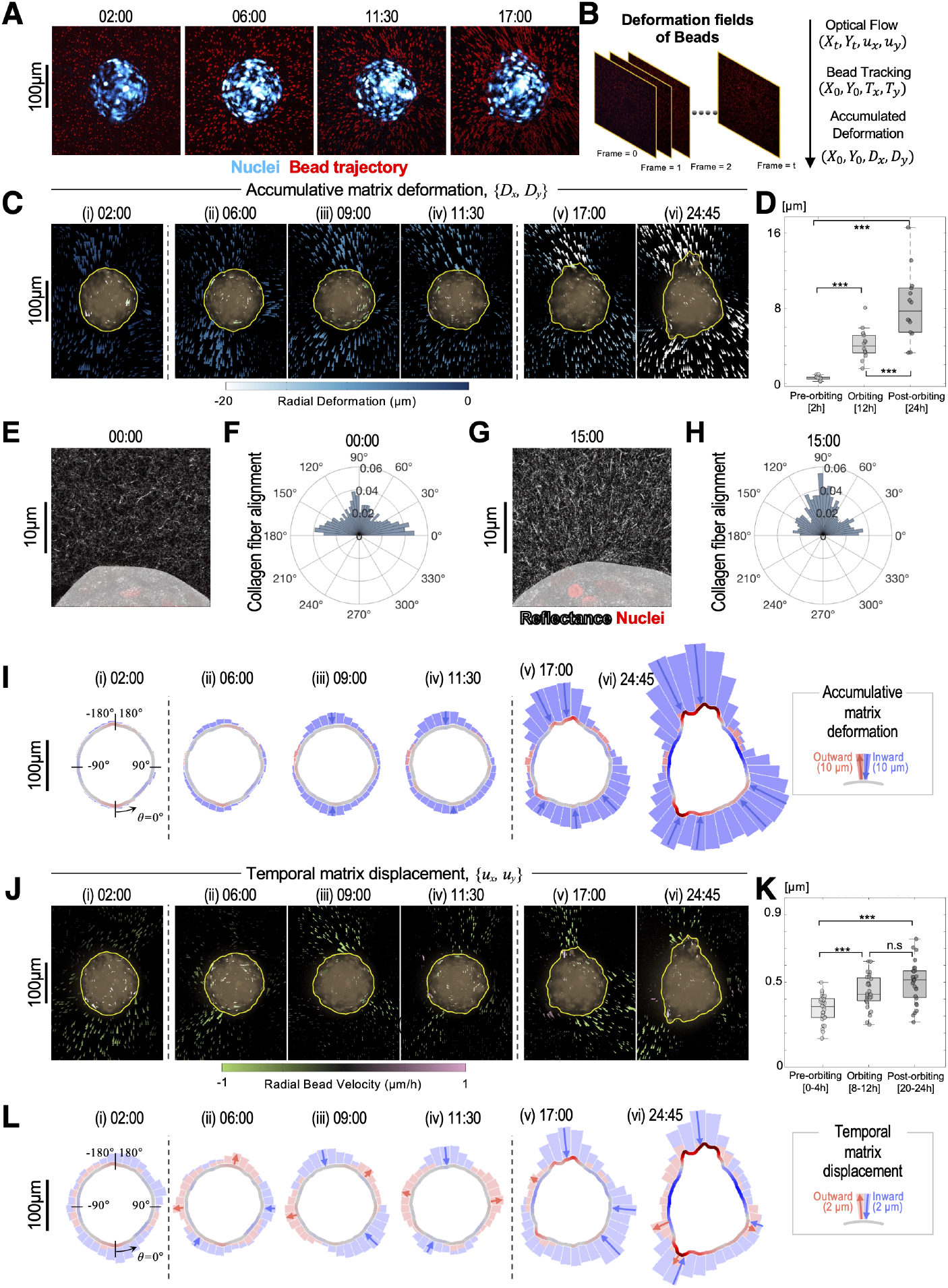
Spheroids exert non-uniform deformations on the surrounding matrix, with hotspots along the initial elongation axis. **(A)** Representative snapshots of fluorescent nuclei (blue) within spheroid and tracer particles (red) in the surrounding matrix. **(B)** Schematic of optical flow-based analysis for accumulative displacement of the tracer particle field corresponding to matrix deformation. **(C)** The accumulative displacement vectors of tracer particles at the different stage of cell migration(i:Pre-orbiting, ii-iv:Orbiting, v,vi:Invading). **(D)** Comparison of accumulative bead displacements based on phase in multiple samples (n=3, spheroids=14). The displacement was quantified using the amplitude of undulation patterns in radial displacement along angular coordinates, repeated-measures of ANOVA was conducted for the statistical analysis. **(E, F)** Confocal reflectance imaging of isotropic collagen fiber orientation immediately after embedding. **(G, H)** Confocal reflectance imaging of aligned collagen fiber orientation immediately after embedding. **(I)** DAR^1^T^1^ visualization of inward/outward cumulative particle displacement at various binned angular positions. **(J)** The temporal displacement vectors of tracer particles in the matrix at the different stage of cell migration (i:Pre-orbiting, ii-iv:Orbiting, v,vi:Invading). Temporal displacements are calculated sequentially from a reference state set 2 h prior to the snapshot shown. **(K)** Comparison of 75th percentile of temporal displacement magnitudes around the spheroids (n=3, spheroids=14) regarding phases, the Kruskal-Wallis H test and Bonferrnoi correction were employed for the statistical analysis. **(L)** DART visualization of inward/outward temporal particle displacement at various binned angular positions.

### Reversible Suppression of Collective Migration by Osmotic Pressure

We then sought to manipulate collective migration by applying osmotic pressure to perturb cellular outgrowths at the spheroid periphery. Cell culture media was supplemented with 4% polyethylene glycol 400 Da (PEG400) to increase the osmolality, which resulted in a volumetric compression (Fig. 4, S8). For example, multicellular spheroids in the control condition with normal (isotonic) media exhibited the usual orbiting for 1 day, followed by a transition to invasion over the next 4 days (Fig. 4ACD, S8AEFG). In comparison, spheroids continuously treated with PEG400 over 5 days maintained a circular morphology with roughly constant size (Fig. 4BEF, S8B). Moreover, cell migration was largely arrested, with minimal coordination and tractions (Fig. S8HIJ, S9A).

**FIG. 4.**
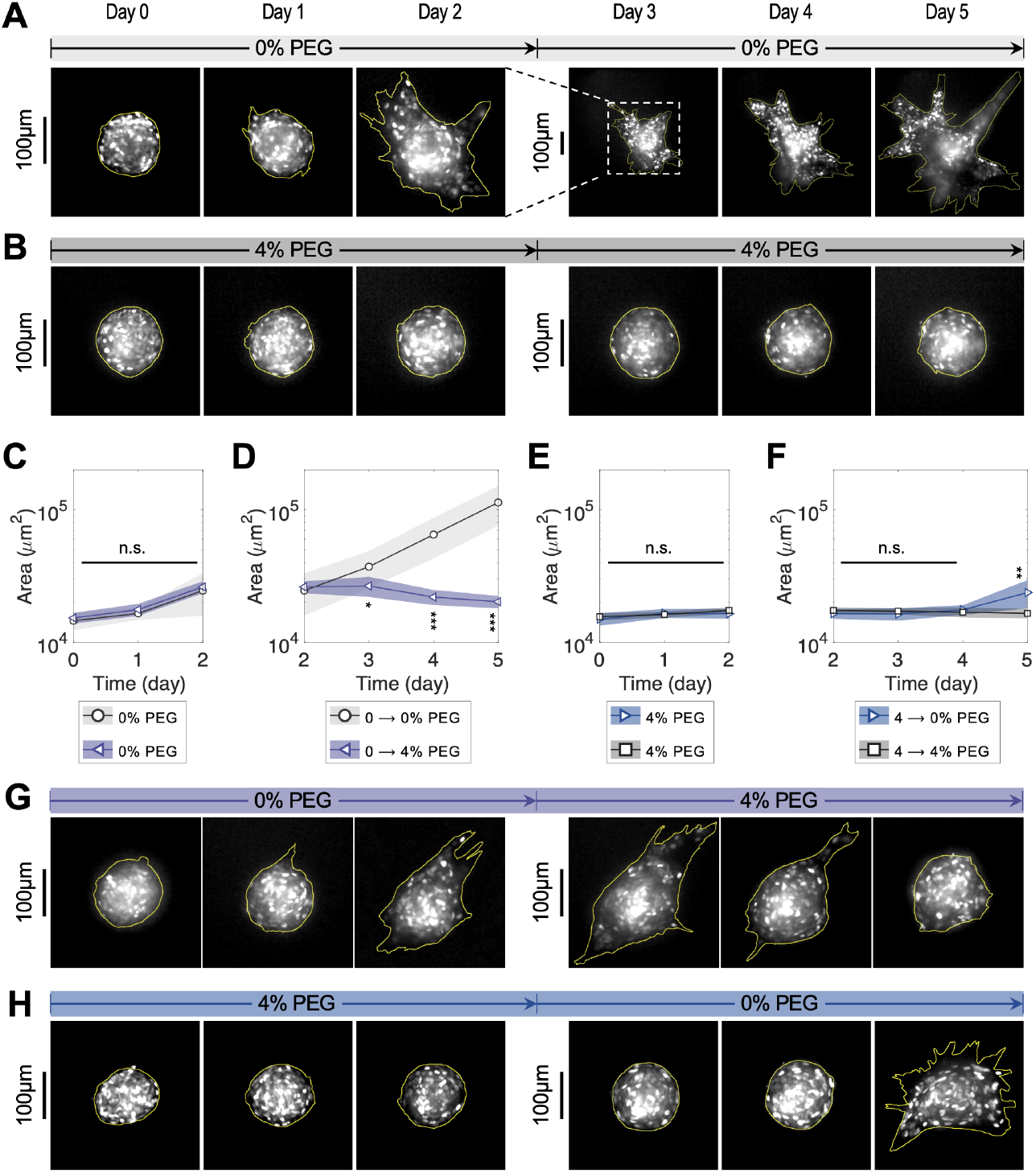
Osmotic compression can transiently arrest collective orbiting or reverse matrix invasion. **(A)** Representative snapshots of coordinated orbiting and invasion over 5 days in normal isotonic media. **(B)** Spheroids are arrested over 5 days in hypertonic media (4% PEG400). **(C-F)** Projected area of the spheroids under each condition. For the first 2 days, Spheroids were maintained in either 0% or 4% PEG media (C, E). Afterwards, selected spheroids were transferred to media with a different osmotic condition (4% or 0%, respectively). Two-tailed t-test, \**p <* 0.05. **(G)** Spheroids initially orbit and invade for 2 days in isotonic media, but then invasion is reversed for the next 3 days in hypertonic media (4% PEG400). **(H)** Spheroids are arrested over 2 days in hypertonic media (4% PEG400), then orbit and invade over the next 3 days in isotonic media.

Further, we investigated whether this suppression of collective migration by osmotic pressure was reversible. First, multicellular spheroids were cultured for 2 days in normal media, exhibiting the usual circumferential orbiting on day 1 and radial invasion on day 2 (Fig. 4G, S8C). With the addition of PEG400-supplemented media, matrix invasion was arrested. Remarkably, the multicellular strands gradually retracted back into the spheroid over the next 2 days, and the spheroid was roughly spherical by day 5 (Fig. 4G, S8C). This switch from normal media to PEG400 after day 2 resulted in a monotonic decrease in projected area and traction, along with increase in circularity, which was the opposite of the control condition (Fig. 4C,D, S9B). It should be noted that these spheroids did not exhibit any orbiting motion at day 5S9KLM), indicating that collective migration was largely arrested by osmotic pressure, consistent with the spheroids continuously treated with PEG400 (Fig. 4B).

Spheroids initially treated with PEG400 remained compact with minimal cell migration over 2 days. After replacement with normal media on day 2, spheroids remained round but increased slightly in size on day 3 (Fig. 4E,F, and H). Nevertheless, orbiting was observed on day 4, representing a delayed onset relative to the control condition (Fig. S8NOP). In accordance with previous results, inward traction increased as the spheroid orbiting motion initiated (Fig. S9D,E). Further, radial invasion was observed on day 5. Thus, this switch from PEG400 to normal media after day 2 led to a slight increase in projected area only on day 5, along with a sizable decrease in circularity. Thus, transient osmotic pressure treatment can delay collective orbiting and even reverse matrix invasion of 3D multicellular spheroids.

### Increasing Heterogeneity Disrupts Collective Orbiting but not Radial Invasion

We next explored whether differential cell-cell adhesion would affect collective orbiting or matrix invasion through the addition of a second cell type (also MCF-10A), which could be induced to undergo the epithelial-mesenchymal transition (via Snail1) to downregulate cell-cell adhesion and upregulate vimentin intermediate filament expression [37]. Fluorescently labeled MCF-10A Snail cells were mixed with MCF-10A wildtype (WT) cells at varying ratios at constant total cell numbers in the hanging drop to generate “mosaic” spheroids, then embedded in collagen matrix as before.

Spheroids with 10% Snail cells exhibited collective orbiting followed by matrix invasion (Fig. 5ABC), comparable to the wholly WT spheroids (Fig. 1AB). Remarkably, both Snail and WT cells at the periphery exhibited coordinated orbiting (Fig. 5D). Subsequently, some Snail cells localized at the tips of invasive strands, suggestive of “leader cells” (Fig. 5E). Matrix displacements were comparable for mosaic spheroids with 10% Snail and wholly WT spheroids (Fig. S10A). In comparison, spheroids with 30% Snail cells did not exhibit collective orbiting (Fig. 5FGHI). 30% Snail spheroids did exhibit several invasive strands, often led by Snail cells (Fig. 5J). Matrix displacements increased continuously over time for 30% Snail mosaic spheroids (Fig. S10B).

**FIG. 5.**
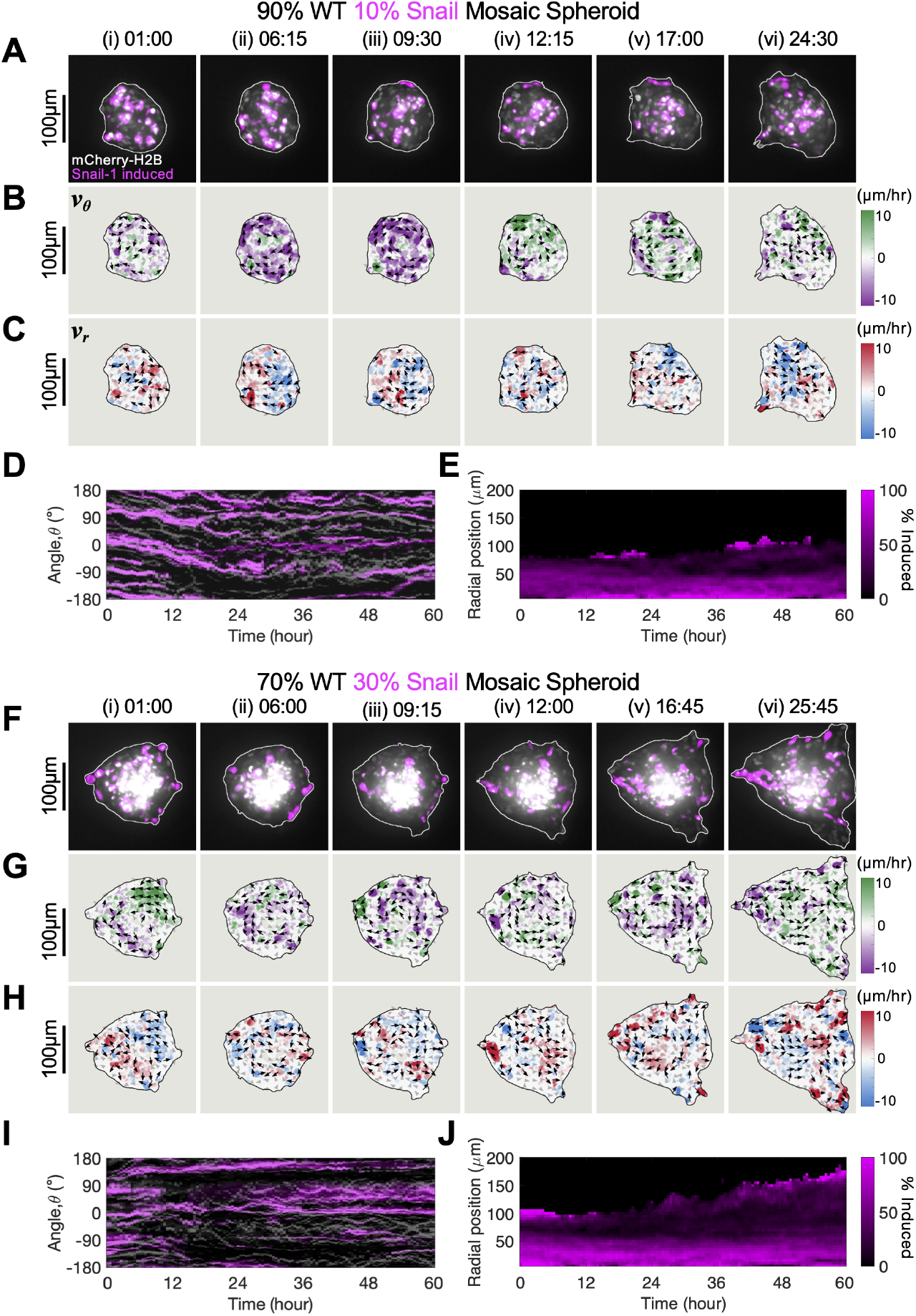
Coordinated migration is impeded by increasing fractions of Snail-induced cells (pink) mixed with wild type (WT) cells in mosaic spheroids. **(A)** Representative snapshots of 10% Snail and 90% WT cells in mosaic spheroids. **(B, C)** spheroids exhibit unimpeded circumferential velocities and hotspots of radial velocity corresponding to matrix invasion. **(D)** Kymograph shows coordinated orbiting of both Snail and WT cells from 0-24 h. **(E)** Snail cells are sorted both the interior and periphery of the spheroid. **(F)** Representative snapshots of 30% Snail and 70% WT cells in mosaic spheroids. **(G, H)** spheroids exhibit weakened circumferential velocities but hotspots of radial velocity corresponding to matrix invasion. **(I)** Kymograph shows no coordinated orbiting of both Snail and WT cells from 0-24 h. **(J)** Snail cells are again sorted both the interior and periphery of the spheroid.

Finally, spheroids composed of 100% Snail cells maintained a rounded morphology over 48 h, with negligible collective migration, either circumferential or radial (Fig. S11) with very weak inward and outward matrix displacements (Fig. S10C). This suggests that individual MCF-10A Snail cells are unable to sufficiently deform and remodel the surrounding collagen I for invasion, perhaps lacking sufficiently strong cell-cell adhesions.

We then treated with a broad spectrum MMP inhibitor (GM6001) compared to DMSO as a vehicle control. 100% WT spheroids after GM6001 treatment exhibited clockwise orbiting from 6-30 h, with a gradual reversal towards counter-clockwise orbiting that continued through 48 h (Fig. S12). These spheroids became more circular after 12 h, but maintained this rounded morphology through the next 5 days. Thus, multicellular spheroids sustain orbiting migration when they cannot easily remodel the surrounding matrix. In comparison, mosaic spheroids of 30% Snail treated with GM6001 exhibited slow matrix invasion with no orbiting (Fig. S12). Altogether, these results demonstrate that collective orbiting is sustained by inhibited matrix remodeling, but is impeded by increasing fractions of Snail cells.

### Cellular Adhesions and Boundary Geometry Govern Coordinated Orbiting in a Computational Model

To further elucidate how collective orbiting was mediated by interactions between cells and the ECM boundary, we developed a computational model with tunable interactions and analyzed how perturbations to this system affected the stability of coordinated migration. Cells were treated as active, self-propelled agents that interact via soft repulsion at short distances and adhesion-based attraction at longer distances [33, 38, 39]. Collective orbiting (“milling”) can emerge in the absence of boundaries when their cell-cell forces are defined by certain potentials [35]. Nevertheless, here we further implement an ECM boundary as a set of discrete equally spaced particles that surround the cells. In order to follow a curved boundary, cells were polarized with an orientation axis and can only adhere to ECM particles within a certain angular range of this axis (± 60^°^). Further, cells exhibit an analogous short-range soft repulsion from ECM particles (see Supporting Methods for further information). Since we aimed to understand how the interplay of cellcell and cell-matrix forces influences collective orbiting, we made the simplifying assumption that all ECM particles were fixed to eliminate potentially confounding effects from dynamic boundaries.

We systematically varied parameters associated with cell-cell and cell-matrix adhesion forces acting on cells confined within a perfect circular boundary to evaluate how these interactions (de)stabilize collective orbiting over long time periods. We considered the average normalized angular momentum ⟨*L*⟩, which has a value of 1 for perfectly coordinated angular motion and 0 for random migration. We found migration was arrested in the limit of weak cell-matrix adhesion and strong cell-cell adhesion (Fig. 6AB, S13). Globally coordinated orbiting occurred in an intermediate regime with slightly stronger cell-matrix adhesion and slightly weaker cell-cell adhesion. Further increases in the cell-matrix adhesion strength parameter resulted in peripheral orbiting of the outermost layers but uncoordinated migration of interior cells (Fig. 6AB).

**FIG. 6.**
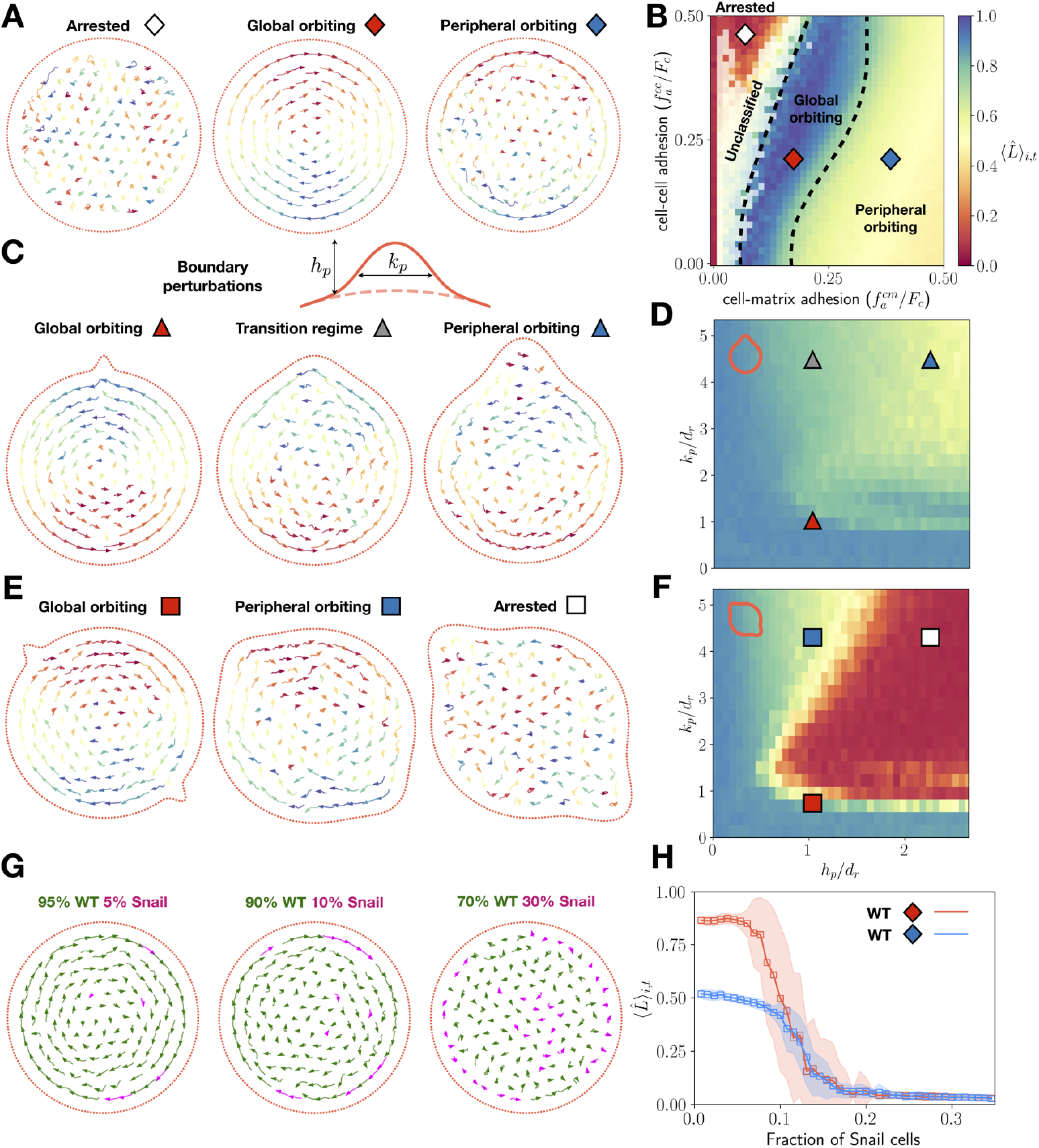
Self-propelled particle model elucidates crosstalk of cell-cell and cell-matrix adhesion with boundary geometry. **(A, B)** Representative simulations of uncoordinated (white diamond), global orbiting (red diamond) and peripheral orbiting (blue diamond) states, and the corresponding phase diagram of the angular order parameter ⟨*L*⟩ over the last 100 hours of the simulation across various cell-cell and cell-matrix adhesion strength parameters. The dashed lines on the phase diagram are hand-drawn and approximate the boundaries of the three states. **(C, D)** Representative snapshots of global (red triangle), transitory (grey triangle) and peripheral orbiting (blue triangle) associated with a single boundary perturbation, and the corresponding phase diagram of the average order parameter across varying perturbation heights (*hp*) and widths (*kp*). Orbiting is generally maintained with a single boundary perturbation, regardless of its width and height, since particles can follow a relatively large continuous section of curved boundary. **(E, F)** Representative snapshots of global orbiting (red square), peripheral orbiting (blue square) and uncoordinated migration (white square) with two oppositely-oriented boundary perturbations, and the corresponding phase diagram depicting how the angular order parameter phase diagram responds to different perturbation heights and widths. Global orbiting transitions towards uncoordinated motility with two boundary perturbations of increasing width and height, due to cells sensing smaller continuous section of curved boundary. **(G, H)** Snapshots of simulations with increasing percentages of Snail+ cells, which are assumed to have lower cell-cell adhesion and higher cell-matrix attraction compared to WT cells. Diamonds indicate the WT adhesion parameters in the phase diagram in (A). The corresponding phase plot of the angular order parameter as a function of Snail percentage demonstrates how global orbiting is disrupted by increasing fractions of Snail cells, which sort to the boundary and impede collective migration.

We then considered the stability of collective orbiting to geometric perturbations of the boundary. Collective orbiting was relatively stable to one outward perturbation, since cells migrated by following the circular boundary and detoured around misaligned cells at the perturbation site (Fig. 6CD). However, two sharp features with height *h*_*p*_ greater than one cell diameter and width *k*_*p*_ greater than two cell diameters were sufficient to destabilize collective orbiting (Fig. 6EF). Cells now encountered multiple obstacles and had shorter lengths of continuously curved boundary to follow. The number and size of these boundary perturbations are consistent with our experimental observations (Fig. S14).

Finally, we analyzed collective orbiting in a mixture of two cell types with distinct adhesion properties. Snail cells were implemented with weaker cell-cell adhesion and stronger cell-matrix adhesions than WT cells, as well as isotropic sensing of the matrix. We observed that collective orbiting was robust to relatively small fractions of Snail cells (up to ∼ 10%), since WT cells at the periphery could orbit by migrating around the Snail cells at the periphery (Fig. 6GH). However, initializing simulations with greater Snail cell fractions caused the cell-matrix interface to become fully occupied by Snail cells, which impeded the ability of WT cells to migrate along the boundary. Overall, these simulations demonstrate the role of boundary curvature for coordinated orbiting with comparable cell-cell and cell-matrix adhesion. Further, an increasing fraction of obstructionist cells that occupy the boundary can also destabilize collective orbiting.

## III. DISCUSSION AND CONCLUSION

Epithelial tissues often transition between spherical to elongated morphologies via coordinated orbiting and matrix remodeling. For example, Drosophila egg chambers elongate via global rotation and basement membrane deposition [9, 10]. Moreover, solid tumors often exhibit ellipsoidal morphologies, which has been attributed to asymmetric stress distribution in a surrounding stiff matrix [40, 41]. Finally, mammary organoids transition from cylindrical branches to spherical alveoli when the local “surface tension” switches from axial to circumferential, stabilized by collective rotation [6]. Our experiments reveal a remarkably deterministic transition from orbiting to matrix invasion driven by the dynamic reciprocity of cells and matrix at curved interfaces.

We implemented a computational model of self-propelled particles with tunable cell-cell and cell-matrix interactions confined within an arbitrary geometry. Our model shows that orbiting is robust when cell-cell and cell-matrix adhesions are comparable, with few sharp boundary perturbations and mostly wildtype cells. The computational results are in agreement with the experimental observations, but we acknowledge several simplifying assumptions. We treat the boundary as fixed and rigid since cell migration occurs on considerably faster timescales than matrix remodeling and proliferation. Our results thus represent a limiting case for finding parameter regimes where coordinated orbiting can occur robustly. Nevertheless, future modeling could incorporate dynamic reciprocity between cells and matrix [42–48], as well as proliferation [49] and the fibrous architecture of the matrix [42, 43, 45, 46].

In conclusion, we show that dynamic reciprocity between multicellular spheroids and the surrounding matrix drive a transition from coordinated orbiting to strand invasion. We observed that spheroids were initially ellipsoidal, but became more rounded during circumferential orbiting. This “flattening” of the spheroid along the major axis corresponded to locally increased matrix displacements, which are subsequently locations of multicellular strand invasion in the radial direction. We show that these orbiting and invasion dynamics can be transiently arrested and reversed through the application of osmotic pressure. Moreover, we demonstrate that both orbiting and invasion can occur in mosaic spheroids with a small fraction of EMT induced cells with weakened cell-cell adhesions. Nevertheless, an increasing fraction of EMT induced cells will impede orbiting but permit matrix invasion. Lastly, we implement a minimal physical model to elucidate the interplay of cell-cell and cell-matrix adhesion with local curvature in maintaining collective orbiting. Ultimately, this spherical symmetry-breaking mechanism via a collective transition from circumferential to radial motility has potential relevance for epithelial tissue morphogenesis towards more topologically complex architectures via local budding and branching.

## ACKNOWLEDGMENTS

We thank S.E. Leggett, R.E. Baker, J. Notbohm, D. Bhaskar, J. Yang, and A. McGhee for helpful conversations, as well as J.S. Brugge and D.A. Haber for the gift of stably transfected MCF-10A cell lines. IYW and JAC also thank P. Kulesa and P. Maini for catalyzing this collaboration. We acknowledge funding from Brown University’s Hibbitt Engineering Postdoctoral Fellowship (JK), NIH R01GM140108 (JK, HJ, AMH, AM, HY, MG, IYW), “la Caixa” Foundation Fellowship 100010434 with code LCF/BQ/EU21/11890128 (C. Falcó), EPSRC grant EP/R014604/1 (C. Falcó, WDM, JAC), ERC Horizon 2020 Research and Innovation Program Advanced Grant Non-local-CPD 883363 (WDM, JAC), ONR Panther Award N000142212828 (MA, C. Franck), MIT School of Engineering Takeda Fellowship (HY), and ARO W911NF2310385 (IYW). WDM and JAC would also like to thank the Isaac Newton Institute for Mathematical Sciences, Cambridge, for support and hospitality during the program “Mathematics of movement: an interdisciplinary approach to mutual challenges in animal ecology and cell biology”, where work on this paper was undertaken.

## DATA AVAILABILITY

All code used to simulate the model and produce the figures in the main text is available on Github.

## AUTHOR CONTRIBUTIONS

IYW conceived and supervised the project. JK and IYW designed experimental work. JK, AMH, and AM performed spheroid experiments. JK, HJ, MA, C. Franck, and IYW analyzed collective migration and tractions. JK, HY and MG characterized matrix architecture. C. Falcó, WDM, JAC and IYW designed computational and theoretical work. C. Falcó and WDM implemented theoretical model and performed simulations. JK, HJ, C. Falcó, WDM, and IYW wrote the manuscript with feedback from all authors.

## MATERIALS AND METHODS

### Cell Culture

Human mammary epithelial cells (MCF-10A), stably transfected with H2B-mCherry as a fluorescent nuclear marker were a generous gift from M.R. Ng and J.S. Brugge (Harvard Medical School). Briefly, MCF-10A cells were grown on standard tissue culture T-75 flasks in a 5% CO_2_ humidified incubator at 37 °C. The growth medium is composed of DMEM/F12 (Invitrogen, 11965-118) supplemented with 5% horse serum (Invitrogen, 16050-122), 20 ng/mL epidermal growth factor (R×D Systems, 236-EG), 10 *µ*g/mL insulin (Sigma Aldrich, I-1882), 0.5 *µ*g/mL hydrocortisone (Sigma Aldrich, H-0888), 100 ng/mL cholera toxin (Sigma Aldrich, C-8052), and penicillin streptomycin (Invitrogen No. 15070-063).

An EMT-inducible MCF-10A variant was stably transfected with an 4-hydroxy-tamoxifen (4-OHT) inducible Snail expression construct fused to an estrogen receptor element response element (ER-Snail-1^6SA^), a generous gift from D. A. Haber (Massachusetts General Hospital)^1^. The Snail-1^6SA^ variant is refractory to phosphorylation and is thus stably expressed and localized in the nucleus, where it initiates EMT induction. This cell line also overexpresses fluorescent proteins in the nucleus (mCherry-H2B) and cytoplasm (GFP) for live cell tracking. 4-OHT resuspended at DMSO were added to the MCF-10A growth media at a final concentration of 500 nM for 72 h to induce Snail induction and EMT.

### Multicellular Spheroid Preparation

Multicellular spheroids were prepared using a modified hanging drop method. Briefly, single cell suspensions of MCF-10A were diluted to 20,000 cells/mL with methycellulose (MethoCel(R) A4M, Sigma Aldrich, 94378) at a final concentration of 1 mg/mL to promote spheroid aggregation. 25 *µ*L droplets (500 cells each) were then dispensed onto Petri dish lids using a multichannel pipettes. These lids were gently flipped over above dishes containing 8 mL of PBS supplemented with 1% penicillin streptomycin. After 48 h, the cells aggregated into multicellular spheroids and could be isolated for further investigations. Most experiments were conducted using MCF-10A-H2B-mCherry cells. Mosaic spheroids were prepared by mixing OHT-treated MCF-10A-ER-Snail-1^6SA^ at 10%, 30% or 100% of total cell count, with the remainder being MCF-10A-H2B-mCherry cells.

### Spheroid Embedding in 3D Collagen I Matrix

Bovine collagen I solution (FibriCol(R), Advanced Biomatrix, 5133) was prepared at a final concentration of 6 mg/mL for spheroid embedding. Briefly, collagen I stock solution was first diluted with de-ionized water containing fluorescent far-red carboxylated polystyrene microparticles (1 *µ*m diameter with 660/690 nm excitation / emission wavelength, Bangs Laboratories, FCFR006). Microparticles were added for a final concentration of 0.05 mg/mL, after being sonicated in a water bath for 30 minutes. The stock solution was adjusted to pH 7.4 using drops of 1N NaOH, which was determined for every batch using a pH meter. Typically, 14–16 *µ*L of 1N NaOH was added per 1 mL of collagen I precursor solution. 4.5% (v/v) of 10× PBS was added so that the spheroids were not in the hypo-/hyperosmotic condition and prevent salting-out during collagen polymerization. 12.5 mM HEPES was added for buffering effect. The neutralized 7.5 mg/mL collagen solution was then incubated for 1 hour at 4^°^C and then mixed with MCF-10A growth media containing the hanging drop spheroids. The final mixture were seeded in 96-well plate and incubated at 37^°^C. After 1 h of incubation, the hydrogel was overlaid with an additional 200 *µ*L of growth media. To apply osmotic pressure on the embedded spheroids, growth media containing 4% wt/vol (100 mOsml^−^1) polyethylene Glycol 400 (TCI Chemicals, N0443) was overlaid. To inhibit matrix metalloproteinase (MMP) activity, growth media contianing 25 *µ*M GM6001 (Millipore Sigma, CC1010) was used.

### Live-cell Confocal Microscopy and Image processing

Spheroids embedded in the collagen matrix were imaged using Nikon Eclipse Ti fluorescence microscope equipped with a motorized stage, direct illumination LED (Thorburn), a multi-channel light source (Lumencor Spectra-X), spinning disk confocal unit (CrestOptics X-Light V2), cMOS camera (Andor Neo), and a 20× Plan Apo opjective (NA 0.75). Cells were maintained in a microscope-mounted incubator (In Vivo Scientific) under controlled conditions of 37 ^°^C, 5% CO_2_, and humidification. NIS Element software was used for automated image acquisition. MCF-10A-H2B-mCherry, MCF-10A-ER-Snail-1^6SA^, and far-red fluorescent microparticles were excited and detected at 555/620 nm, 395/535 nm, and 640/670 nm, respectively. Z-stack images were acquired every 15 or 30 minutes, capturing at least 5 slices with a z-step size ranging from 0.9-5 *µ*m. Only spheroids whose z-positions were higher than 100 *µ*m above the plate bottom were included for imaging to avoid undesirable 2D migration along the plate substrate. Both the number of z-slices and the z-step size were optimized to maximize the number of spheroids imaged in a single experiment while ensuring high-content imaging quality. Each channel of the z-stacks was processed using maximum intensity projection in ImageJ. Microparticle images captured with 640/670 nm channel were stabilized using ‘Image Stabilizer’ plugin, with the option ‘Log Transformation Coefficients’ option enabled. The transformation log was subsequently applied to the other channels using the ‘Image Stabilizer Log Applier’ plugin, to ensure consistent stabilization across all the channel images. Spheroid binary masks were generated from the stabilized mCherry signal images dictated with 620 nm channel using a custom ImageJ macro. Briefly, unevenly illuminated background was corrected using ‘Substract Background’ function with a rolling ball radius of 50 pixels. Median filter with radius of 4.0 px was applied for denoising. ‘Smooth’ functions was applied, if necessary. The denoised images were binarized with ‘Threshold’ function. To include the nuclei seprated from the center mass, a 30 pixel-wide band was created using ‘Make Band’ function, followed by ‘Enlarge’ with -27 pixels. The completed mask images were saved as image sequences. The stabilized images and mask were later used for kymograph construction and optical flow analysis.

### Kymograph analysis

Kymographs were generated using custom MATLAB code. At each time point, the outermost 20 *µ*m thick shell along the perimeter was selected from the binary mask. The selected region was then sliced at 0.5^°^ intervals. Each slice was superimposed onto the corresponding mCherry signal images. The resulting trapezoid-shaped image slices were interpolated into rectangular shape and then vertically concatenated into a column. The reference angle *θ* = 0^°^ was assigned to the angular position of the point farthest from the centroid in the initial image. Each column from successive time points was again horizontally concatenated to construct the final kymograph. For mosaic spheroids, the same procedure was applied to the EGFP channel image, which was then merged with the kymograph generated from the mCherry signal image.

### Optical flow and angular order parameter analysis

The velocity profiles of cellular migration and bead displacement were obtained using Farneback optical flow algorithm. We adapted ‘opticalFlowFarneback’ function that is available from ‘Computer Vision Toolbox’ of MATLAB, with default property values. For cellular migration analysis, velocity bias due to the abrupt global movement of the spheroid was calibrated by subtracting the mode value of *v*_*x*_ and *v*_*y*_. The resulting *v*_*x*_ and *v*_*y*_ were then converted into *v*_*r*_ = **e**_**r**_ · (*v*_*x*_, *v*_*y*_) and *v*_*θ*_ = **e**_*θ*_ · (*v*_*x*_, *v*_*y*_) where 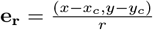 and 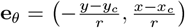, with respect to the centroid of the corresponding binary masks, (*x*_*c*_, *y*_*c*_). Angular order parameter (AOP) was defined to describe the fraction of velocity pixels oriented circumferentially in the outermost region. Outermost 30 *µ*m thick shell along the perimeter was selected as the analysis ROI to sufficiently encompass at least a single cell layer. For each pixel in the ROI, 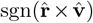 was calculated, where 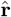 is a radial unit vector from the spheroid centroid and 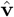 is a unit velocity vector. AOP has a value 0 for a pixel whose velocity is perfectly radially oriented, and 1 or -1 for a pixel whose velocity has a circumferential component. The mean ⟨AOP⟩ was then calculated across all pixels within the analysis ROI. The orbiting phase was defined as the period during which the |⟨AOP⟩| is greater than 0.5, indicating more than 75% of the cells in the ROI are orbiting in the same direction. After identifying the primary orbiting direction (clockwise or counterclockwise), ⟨AOP⟩ during the orbiting phase was adjusted to be always positive, to ensure that the orbiting phase is consistently represented as a crest in the plot.

### ECM pore size estimation

The fibrillar microstructure of collagen I hydrogel was visualized using confocal reflectance imaging. The pore sizes of ECM were estimated based on our previously established method for fiber counting.^2^ Firstly, individual fibers were detected with CT-FIRE. Parameters were kept default except for ‘thresh im2’ and ‘s xlinkbox’ to be manually optimized to be 90 and 2 respectively. Subsequently, fiber features were obtained with CurveAlign. ^3^ Each reflectance image was subdivided into square boxes that are sized 4^2^, 8^2^, 16^2^, 32^2^, 64^2^, 128^2^, 256^2^, 512^2^, and 1024^2^, pixels. The mean number of fibers over all boxes was recorded at various box sizes. A relationship between the box size (box area in *µ*m^2^) and number of fibers was established by constructing a cubic interpolant. To estimate fiber density (and the mesh area) in the image, the area corresponding to a single fiber was approximated using the cubic interpolation function. The pore diameter was then calculated by dividing by *π*, taking the square root and multiplying by 2.

### ECM deformation analysis

Deformation fields in the surrounding extracellular matrix were analyzed by measuring the displacement of tracers using optical flow, which were verified using manual tracking as well as q-factor-based digital image correlation (qDIC^4^). Briefly, far-red fluorescent microparticles with 1 *µ*m diameter were embedded in the collagen I matrix and imaged using spinning disk confocal microscopy using z-slices at 5 *µ*m interval. Since most of the matrix displacements occurred in the x-y plane along the equator, we chose to analyze maximum intensity projections from 9 z-slices centered at the spheroid equator. The embedded beads exhibited continuous trajectories with significant displacement (*>*30*µ*m) over time, making it challenging to correlate images captured at longer time intervals (Fig. 3A, S4A). To track the large displacements over time, we determined the positional changes of the initial grid points (**T**_*grid*_(*t*_0_)) by sequentially accumulating the short-term displacements (**u**_*grid*_(*t*)) between consecutive images(time=*t* − 1 and *t*) captured at 15-20 minutes by adopting the Farneback’s bead movement (Fig. S4B,C,and D). Here, the initial grids for tracking the ECM displacement were generated with a 3.25*µ*m interval in both the x and y directions. The displacement for each updated grid point (**T**_*grid*_(*t*)) at a given time was obtained by interpolating the optical flow displacement field (**u**_*grid*_(*t*)) at its corresponding position akin to the iterative image deformation method built into qDIC. Within this framework, the time interval was sufficiently short to accurately track bead movement (*<*2 µm/h), and the beads exhibited affine trajectories consistent with neighboring beads during this period, confirming the validity of the interpolations.Finally, we mapped the cumulative large displacements(**D**_*grid*_(*t*)) on the Cartesian grids by calculating differences between the updated trajectories(**T**_*grid*_(*t*)) and their initial position(**T**_*grid*_(*t*_0_)) (Fig. S4D and E).

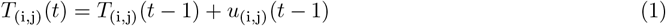

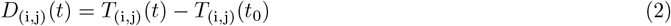

To validate the algorithm for tracking ECM displacement, we compared the calculated cumulative displacement with manually tracked beads. The Pearson correlation coefficients for 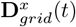 and 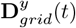 were 0.90 and 0.94, respectively, demonstrating that the algorithm accurately captures the bead movement. Furthermore, the correlation coefficients between qDIC-based and optical flow-based displacement were 0.92 and 0.96, respectively, demonstrating a strong agreement and confirming the reliability of our method for accurately measuring bead displacements (Fig. S4F). All the data interpolations were performed using a cubic spline interpolation algorithm implemented in MATLAB (R2024b, MathWorks, Inc.).

Displacement Arrays of Rendered Tractions (DART) analysis was performed on the 2D displacement data obtained from Optical Flow of bead images. Similar to our previous work,^5^ we constructed a displacement vector field *U*_grid_, where *U*_grid_ is either equal to *D*_grid_ or *u*_grid_, depending on whether DART analysis was performed on the cumulative (*D*_grid_) or incremental (*u*_grid_) displacement field. *U*_grid_ was then decomposed into its radial,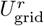, and circumferential, 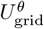, components.

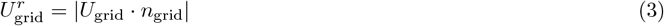

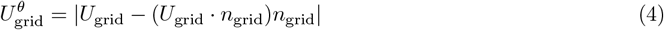

Subsequent classification of each displacement vector into protrusive and contractile was determined by the sign of 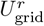. Each particle’s position was binned into one of 36 angular regions depending on their location with respect to the center of the spheroid, and their displacement magnitudes classified according to their value. The mean of each displacement type at each angular region was plotted to obtain a radial DART diagram.

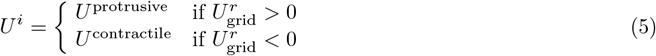

### Information for statistical analysis of deformation and displacement

To quantify the enhancement of deformation patterns associated with the invasive morphology of spheroids, we measured the amplitude of these patterns over time. This was achieved by calculating the average radial deformation in 36 angular sections around the spheroids. A repeated-measures analysis of variance (ANOVA) was then conducted to evaluate the effect of time on the progression of deformation patterns. Specifically, we compared cumulative deformation changes at three key phases: the pre-orbiting phase (2 hours after seeding), the orbiting phase (12 hours after seeding), and the invading phase (24 hours after seeding). Pairwise comparisons were performed using Estimated Marginal Means to assess differences between time points, with Type I error in multiple comparisons corrected using the Bonferroni method. To track changes in temporal displacement magnitude during the orbiting phase, we calculated the 75th percentile of displacement magnitudes around the spheroids, as this metric accounts for the highly localized nature of displacement vectors. The Kruskal-Wallis H test, a non-parametric alternative to one-way ANOVA, was employed to evaluate temporal differences in displacement magnitude. The Bonferroni correction was applied to adjust for multiple comparisons. All statistical analyses were conducted using SPSS (IBM).

### Computational Agent-Based Model

We adapt an existing mathematical model known to produce stable orbiting behavior in the absence of boundaries.^6^ We model cells as spherically symmetric self-propulsive (active) particles. As a first approximation, we take the active force to be proportional to the cell velocity. Cells also experience drag-like forces which act against the cell velocity and is proportional in magnitude to the cube of its speed. The balance between active and drag forces predicts an equilibrium velocity that can be adjusted to match experimental observations. Cells interact with other cells and with the ECM via soft repulsion at short distances and adhesion-based attraction at longer scales. The position of cell *i* at time *t*, **x**_*i*_(*t*), and its velocity, **v**_*i*_(*t*), are calculated using Newton’s second law. The matrix interface is modeled as static discrete particles, with the position of the *k*th particle denoted by **y**_*k*_. The resulting system of ordinary differential equations is given by

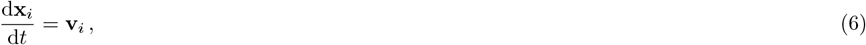

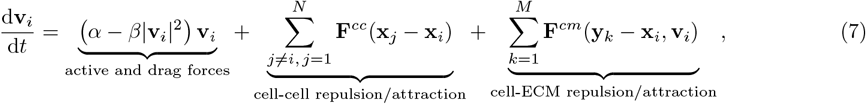

where we have absorbed the mass of each cell into the terms on the right-hand side. We note that previous versions of this model have not considered the influence of the ECM (i.e., the third term on the right hand side of Eqn. (7)) in stabilizing orbiting behavior.

The particular cell-cell and cell-ECM forces that we use are based on those from previous works.^7^ In brief, they ensure that cells repel each other along the direction of contact at short distances, but attract each other at larger scales of interest. They are reproduced in detail in the Supplementary Information (SI), along with details of how we determine parameter values for this framework. Eqs. (6)-(7) are solved with an explicit fourth order Runge-Kutta scheme. Simulations are conducted using the SciPy library in Python. Code to simulate the model is available on Github (see Data Availability). Fifty simulations per parameter regime were used to generate Fig. 6 in the main text. We refer readers interested in more detail about the mathematical model and its implementation to the SI.

### Phase diagrams

We quantify orbiting in numerical simulations using the normalized average angular momentum, 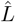, defined as:

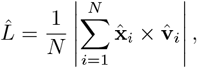

where *N* is the total number of agents and 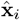 and 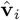 are the unit position and velocity vectors of cell *i*, assuming the system’s center of mass is at the origin. The normalized angular momentum ranges from 0 (no collective coordination) to 1 (perfect orbiting). To construct phase diagrams, we vary cell-cell and cell-matrix adhesion strengths within the interval [0, 0.5*F*_*c*_], where *F*_*c*_ is the maximum adhesive force (see SI for more details). The parameter space is discretized into a 40 × 40 grid, and fifty simulations are performed for each parameter set. We then average 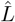 over the last 100 h of each simulation and across all runs. To quantify the time to reach the orbiting state, *τ*, we fit the averaged angular momentum as a function of time, to a sigmoid function with a time scale saturation *τ* (see SI for more details).

### Boundary perturbations

The mathematical model assumes a circular matrix boundary with uniformly distributed particles. To study the impact of boundary geometry we introduce a fixed and static boundary perturbation of height *h*_*p*_ and width *k*_*p*_ by parametrizing the perturbed boundary as

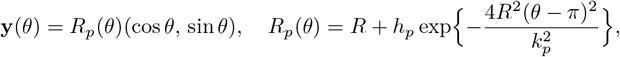

with the perturbation centered at *θ* = *π*. For multiple perturbations the procedure is analogous. We also reparametrize the boundary by arc length parameter to maintain a fixed particle density. To keep a constant cell density, the total number of cells is adjusted according to the total area change induced by the perturbation (see SI for more details).

## Footnotes

1 Wong, I.Y. et al. *Nat. Mater* 13(11), 1063-1071 (2014)

2 Khoo, A.S. et al. *ACS Biomat. Sci. Eng*. 5(9), 4341-4354 (2019).

3 Liu, Y, et al. *Fibrosis: methods and protocols*. 429-451 (2017).

4 Landauer, A. K., et al. *Experimental Mechanics*. 58, 815-830 (2018).

5 Leggett, S.E., et al. *PNAS*. 117(11), 5655-5663 (2020)

6 D’Orsogna, M.R., et al. *Phys Rev Lett*. 96(10), 104302 (2006); Bertozzi, A.L., et al. *Commun Math Sci*. 13(4), 955-985 (2015); Albi, G., et al. *SIAM J. Appl. Math*. 74(3), 794-818 (2014).

7 Carrillo, JA, et al. J. Theor. Biol. 445, 75-91 (2018)

