## Supporting Information for "Collective Transitions from Orbiting to Matrix Invasion in 3D Multicellular Spheroids"

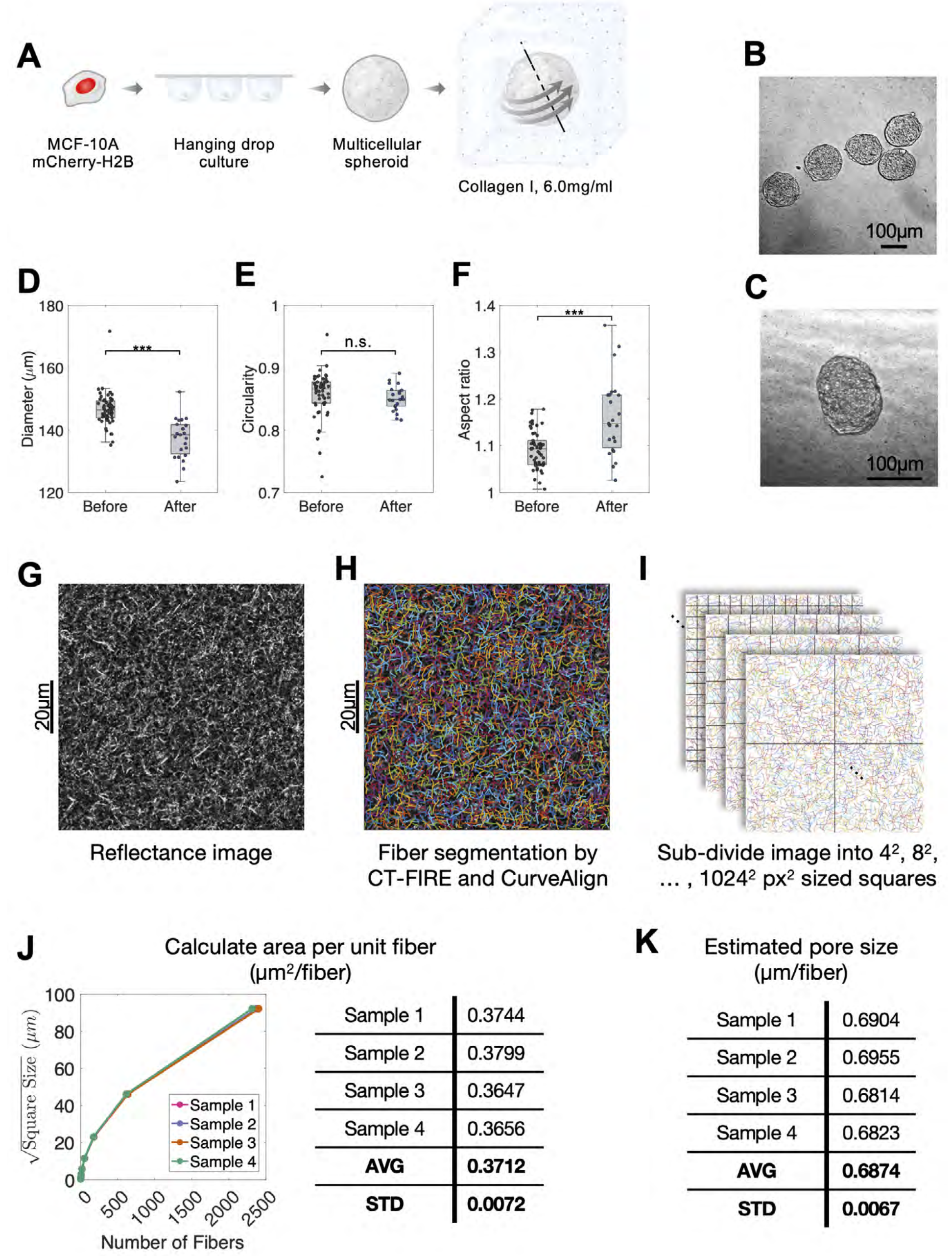


FIG. S1. **(A)** Experimental approach to prepare multicellular spheroids using hanging drops and then embed in collagen I matrix. **(B)** Representative phase image of multicellular spheroids harvested from hanging drop culture. **(c)** Representative phase image of multicellular spheroid embedded in collagen gel. **(D,E,F)** Diameter, circularity and aspect ratio of multicellular spheroids, before and after collagen embedding. Before, 54 spheroids across 4 experiments. After, 34 spheroids across 7 experiments. Two-tailed t-test, $*p < 0.05$. **(G)** Representative confocal reflectance image of collagen I. **(H)** Representative image of fiber segmetation using CT-FIRE and CurveAlign. **(I)** Sub-dividing fiber image for estimation of fiber density. **(J)** Relationship between the number of fiber and the size of square boxes obtained by cubic interpolation. **(K)** Estimated pore size of 4 representative samples.

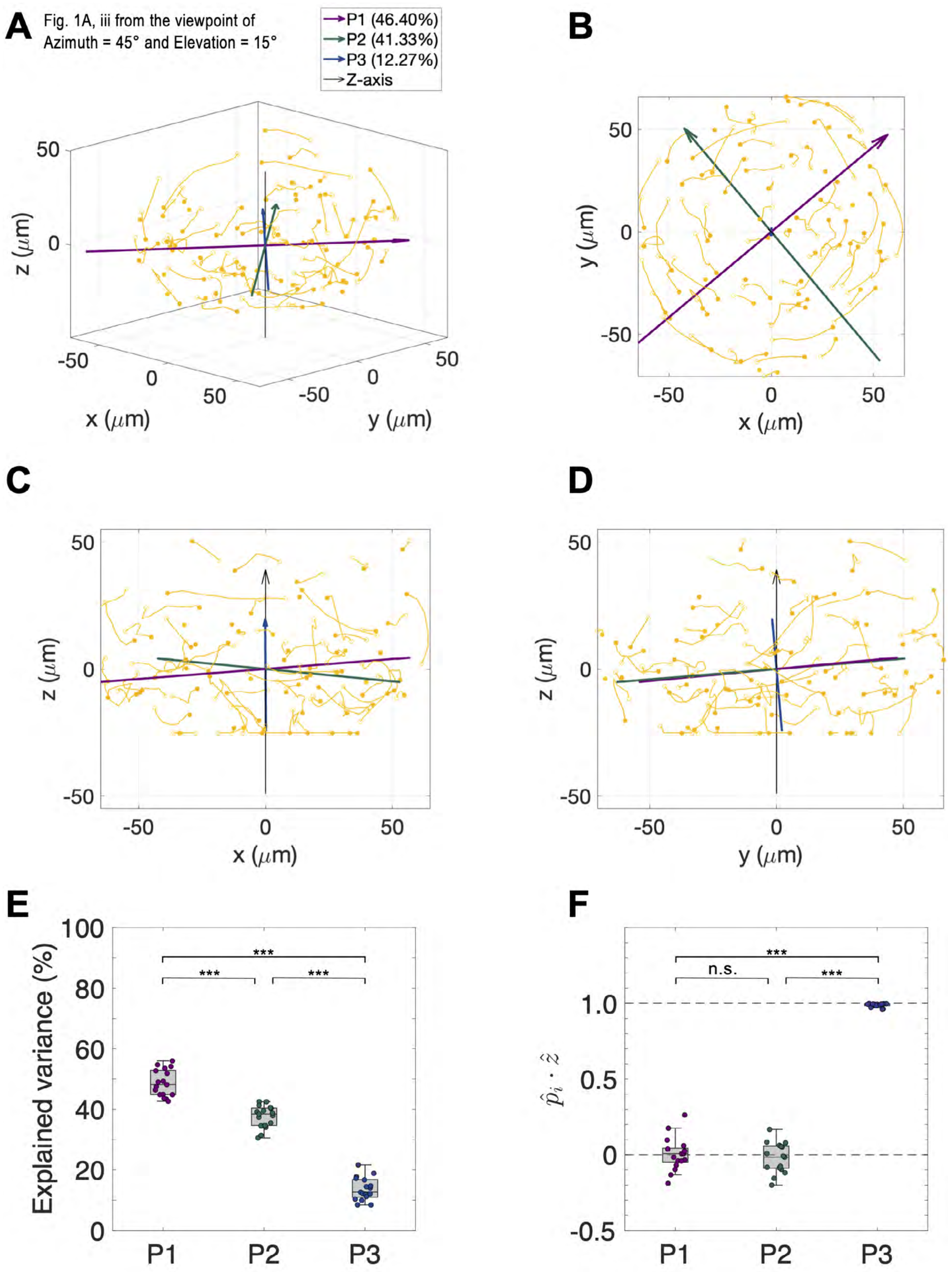


FIG. S2. **(A)** Representative cellular trajectory of Fig. 1A (iii), shown from the viewpoint of Azimuth = 45° and Elevation = 15°. P1, P2, and P3 represent the 3 principal components obtained by performing PCA on the cell trajectory coordinates between 06:00-09:00. **(B-D)** Cellular trajectory and principal components projected on X-Y, Z-X, and Y-Z planes, respectively. P3, the least significant axis was nearly aligned with the z-axis. **(E)** Explained variance of P1, P2, and P3 from 17 spheroids across 2 experiments. P3 accounted for $14 \pm 4\%$ of explained variance.**(F)** Inner product between the 3 principal components and the unit z-vector. Mean $\hat{p}_3 \cdot \hat{z}$ was 0.9895 with standard deviation of 0.0123, confirming the low explanatory power of z-axis. One-way ANOVA with HSD *post hoc,* *$^*p < 0.05$.*

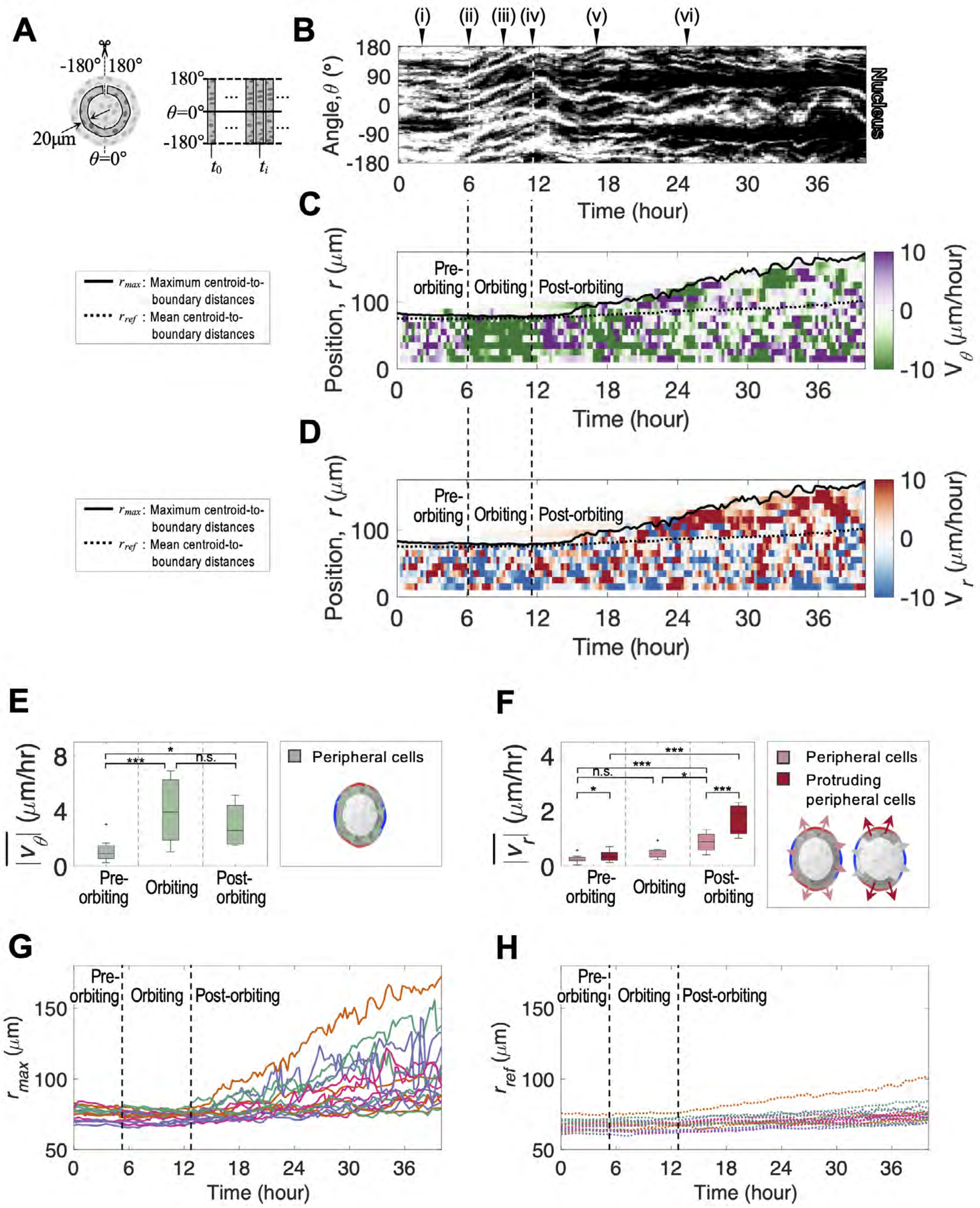


FIG. S3. **(A)** Schematic of unwrapping the next outermost 20 $\mu$m "layer" of the spheroid equatorial plane at each time to generate a kymograph. **(B)** Kymograph of fluorescent nuclei (H2B-mCherry) in the second outermost peripheral layer of cells shows minimal angular motion from 0-6 h after embedding (i-ii). Next, all nuclei exhibit coordinated angular motion from 6-12 h, indicated by the parallel upward trajectories (iii-iv). **(C)** Circumferential velocity from 6-12 h occurs throughout the radius of the spheroid cross-section. **(D)** Radial velocity after 12 h tend to be outwards and large, especially at the spheroid periphery. **(E, F)** Average circumferential velocity during orbiting is significantly larger than before or after, while average radial velocity is faster near protruding poles after orbiting. 8 spheroids across 3 experiments. One-way ANOVA with HSD post hoc, $*p < 0.05$. **(G, H)** Maximum and mean centroid-to-boundary distance over time. Spheroid sizes remain roughly consistent from 0-12 h before increasing during invasion from 12 h onwards.

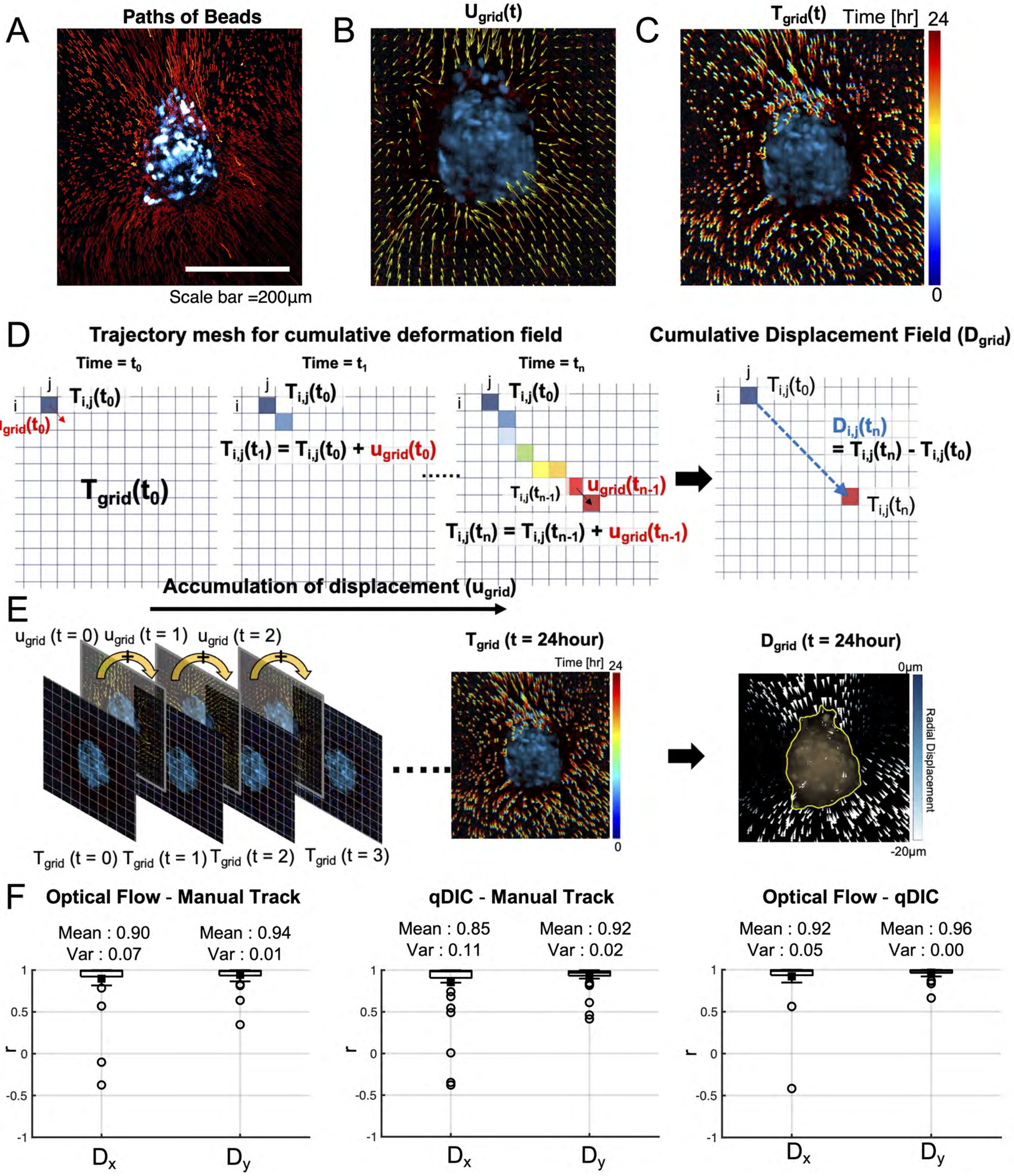


FIG. S4. **(A)** Image for the paths of beads (red) with the cell nucleus (blue) in spheroids, the path was obtained by maximum intensity stacking of beads image for 0-24 hours, **(B)** The results of bead displacement fields from the optical flow calculation,**(C)** The results of trajectories in the ECM grid over time, **(D)** Schematic of updating methods for the trajectory grid and calculation of cumulative displacement field, **(E)** Representative examples for calculating the trajectory grid and cumulative displacement, **(F)** The Pearson correlation coefficient(r) between the two different displacement calculation methods(qDIC and Optical flow) and manual tracking. The correlation was calculated from the temporal arrays of $\mathbf{D}_x(t)$ and $\mathbf{D}_y(t)$ across 4 samples, with a total 40 data points.

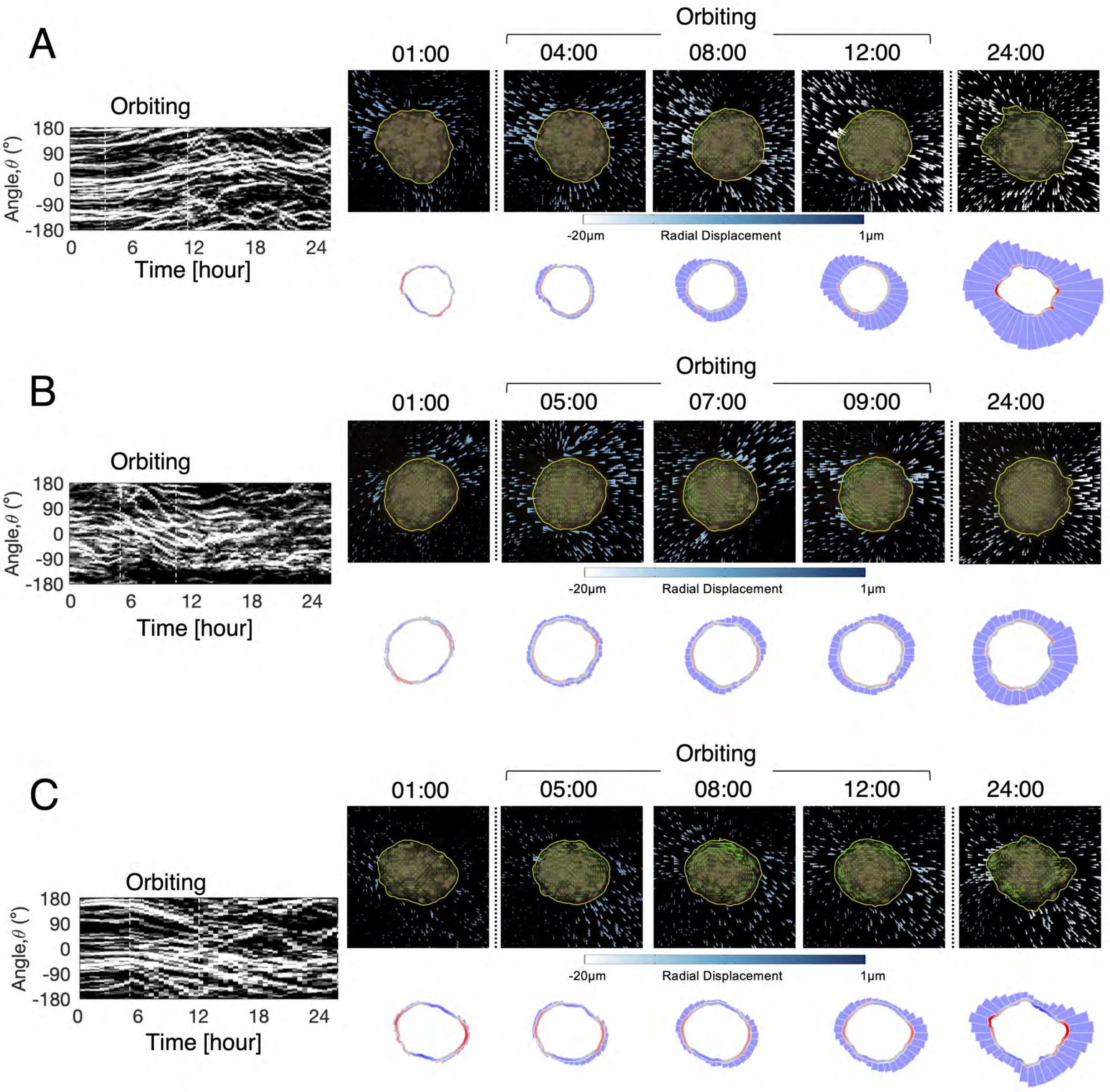


FIG. S5. **(A,B, and C)** Representative kymograph, cumulative bead displacement, and DART plots of WT spheroids. All kymograph shows coordinated circumferential migration patterns during orbiting phases. Bead displacement and DART plots shows asymmetric ECM remodeling during the orbiting spheroids.

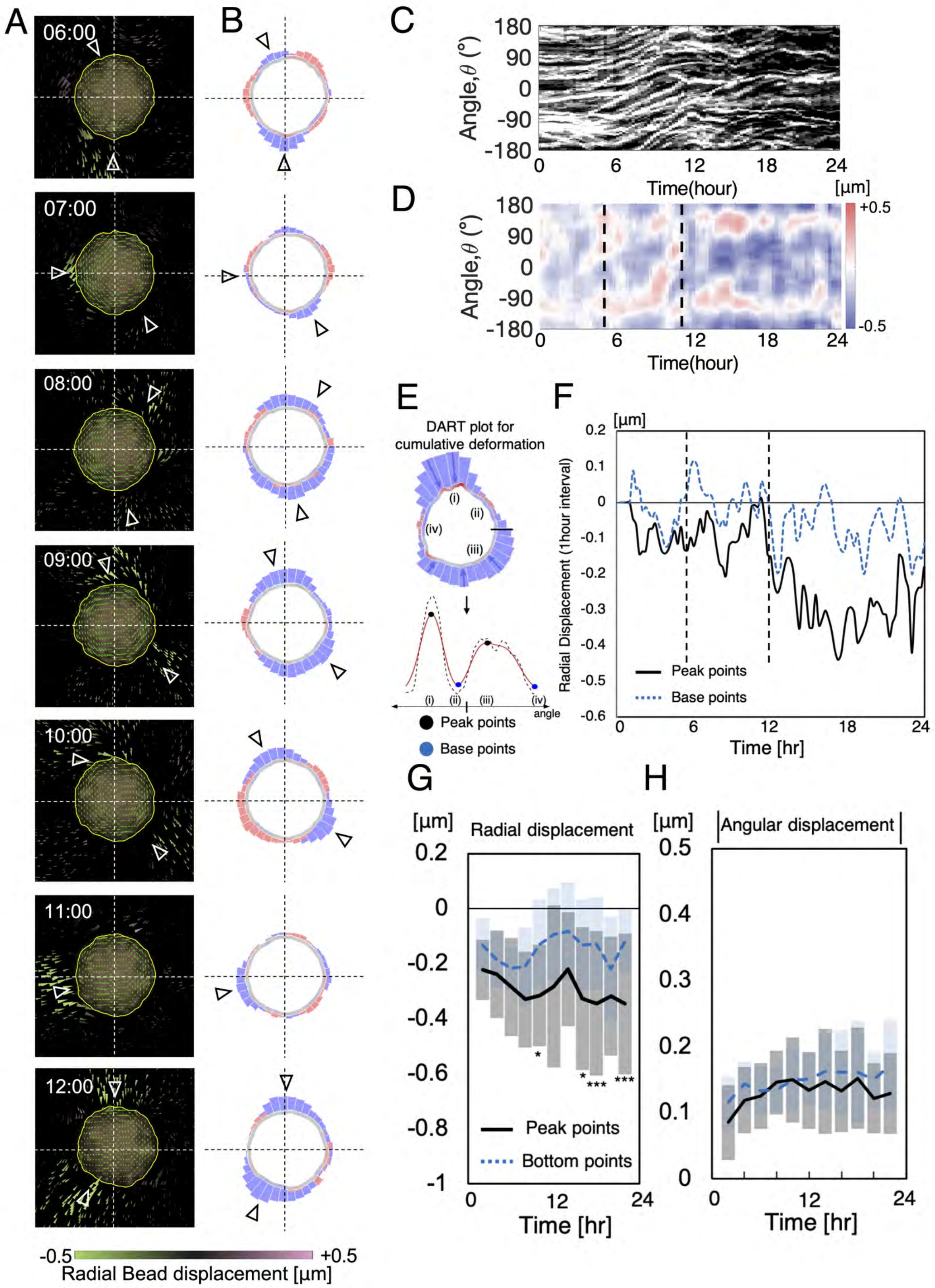


FIG. S6. **(A)** Sequential quiver plots for the temporal displacement of cells over 1-hour intervals, with colors indicating the radial displacement value (green line: the patterns for cell trajectories during the time interval, yellow line: boundary of the spheroid, and white triangle: indicator for the regions where the inward radial displacement are formed), **(B)** Sequential changes in DART plots of radial deformation within the 24 angular binned region, **(C)** Kymographs for the orbiting motion of cells in the spheroids, **(D)** Kymographs for the temporal displacement with colors for radial component value (dotted line: domain for the orbiting time), **(E)** Schematic for analyzing the local changes in radial displacement near the peak and base points of the cumulative radial deformation, **(F)** The changes in radial displacement near the peak and base points, **(G)** Temporal changes in radial displacement analyzed at peak and base points across various conditions (n=3, samples=14), the statistical analysis was conducted by paired-sample t test, **(H)** Temporal changes in the magnitude of angular displacement over time

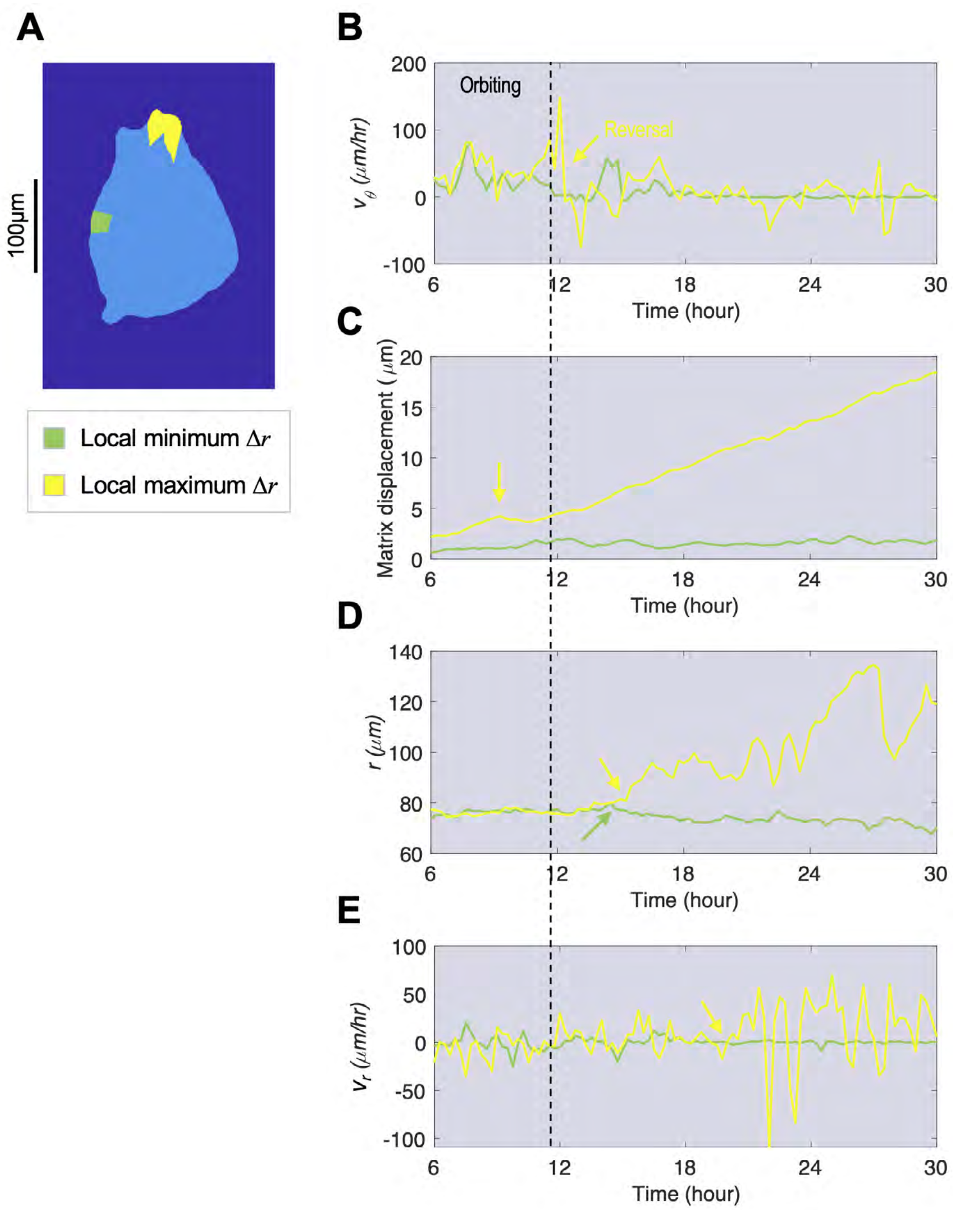


FIG. S7. **(A)** Representation of ROIs with local minimum (green) and local maximum $\Delta r$ (yellow). **(B)** Circumferential velocities of the two ROIs over time. After orbiting phase concludes, $v_\theta$ of ROI with local maximum $\Delta r$ fluctuates over time while gradually damping. ROI with local minimum $\Delta r$ damps sooner. **(C)** ROI with local maximum $\Delta r$ features a temporal peak in matrix displacement at around 9 h, during the orbiting phase. **(D)** The centroid-boundary-distance, $r$, of ROI with local maximum $\Delta r$ abruptly increases at 15 h, while that of ROI with local minimum $\Delta r$ gradually decreases. **(E)** The radial component of cellular velocity exhibit stronger fluctuations from 20 h, making even higher peaks in centroid-boundary-distance.

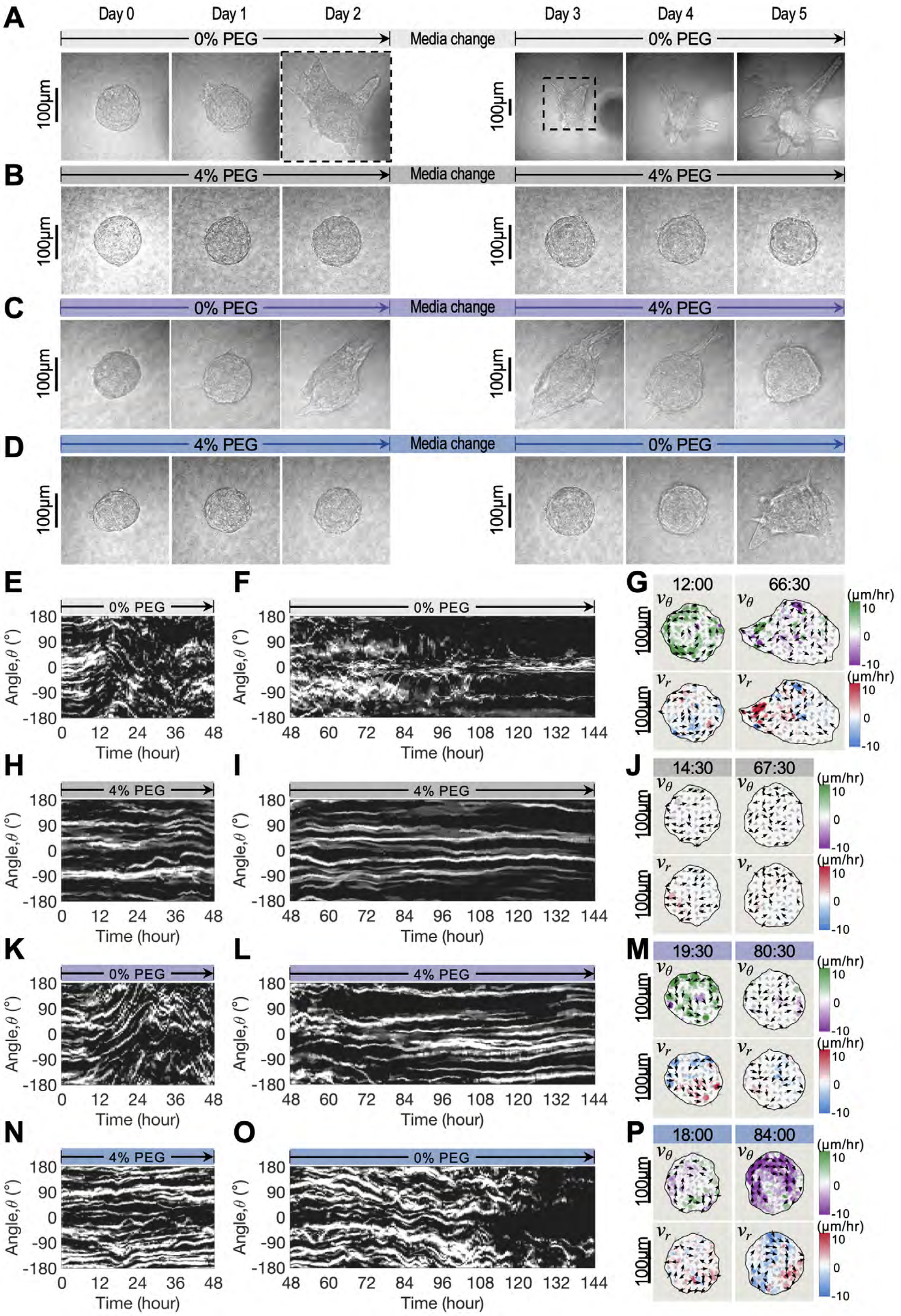


FIG. S8. **(A-D)** Corresponding phase images of Fig. 4 ABGH. **(E,F)** Kymographs for continuous isotonic condition ($0 \rightarrow 0\%$ PEG). **(G)** Representative velocity profiles for continuous isotonic condition. **(H,I)** Kymographs for continuous osmotic condition ($4 \rightarrow 4\%$ PEG). **(J)** Representative velocity profiles for continuous osmotic condition. **(K,L)** Kymographs for isotonic to osmotic condition ($0 \rightarrow 4\%$ PEG). **(M)** Representative velocity profiles for isotonic to osmotic condition. **(N,O)** Kymographs for osmotic to isotonic condition ($4 \rightarrow 0\%$ PEG). **(P)** Representative velocity profiles for osmotic to isotonic condition.

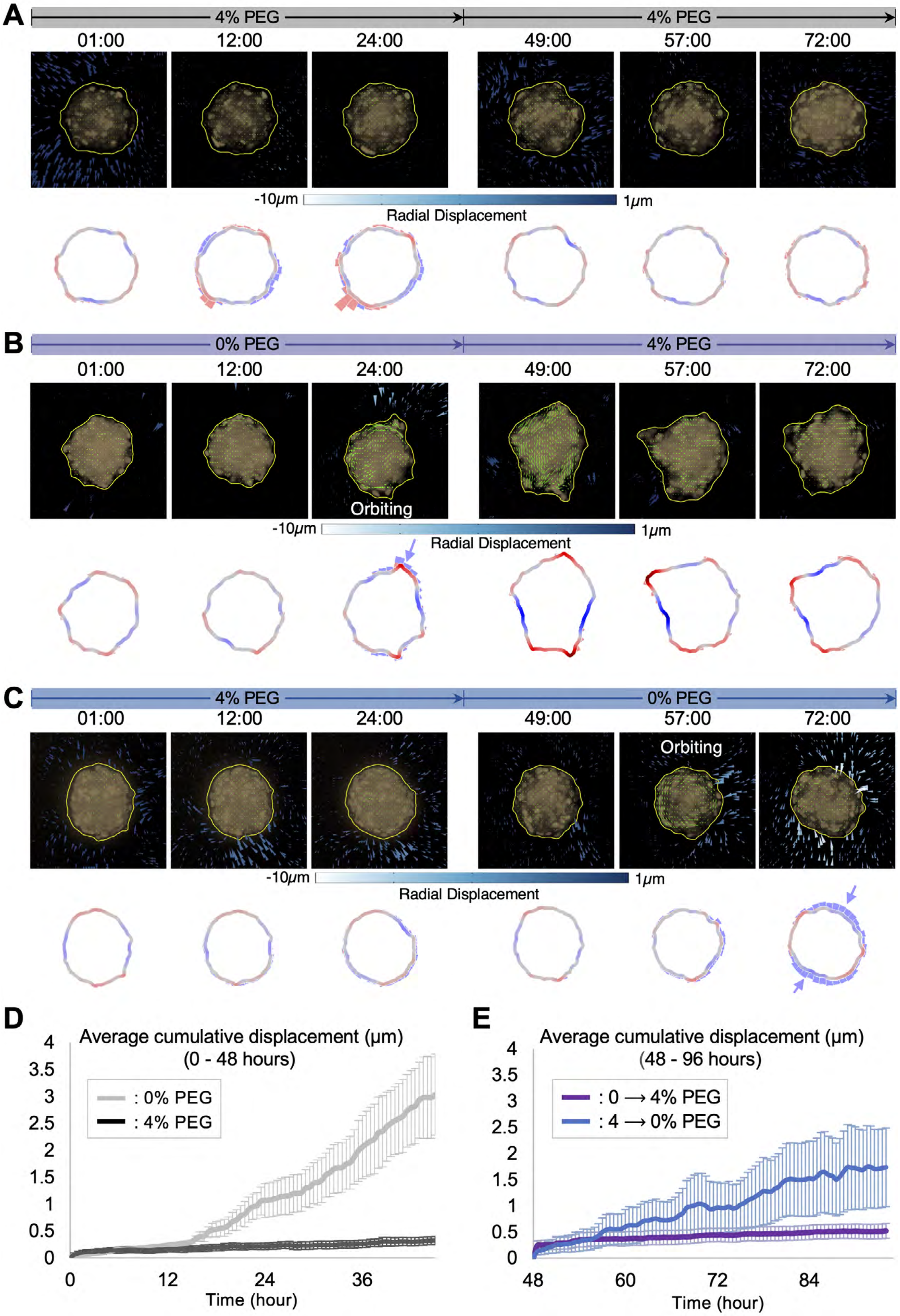


FIG. S9. **(A)** Cumulative bead displacement and DART plot over time with the continuous osmotic condition, 4% PEG, showing no significant contractile displacement of the surrounding ECM. **(B)** Under 0 → 4% PEG condition, the radial contractile displacement disappears shortly after the media change. **(C)** In 4 → 0% PEG condition, spheroids restored their contractility. **(D and E)** The changes in average cumulative displacement near the spheroid according to the osmotic condition. The spheroids under 4 → 0% condition restored their contractility but at a slower and weaker rate compared to those under the unperturbed 0% PEG condition. Error bar: S.E., 20 spheroids across 2 experiments.

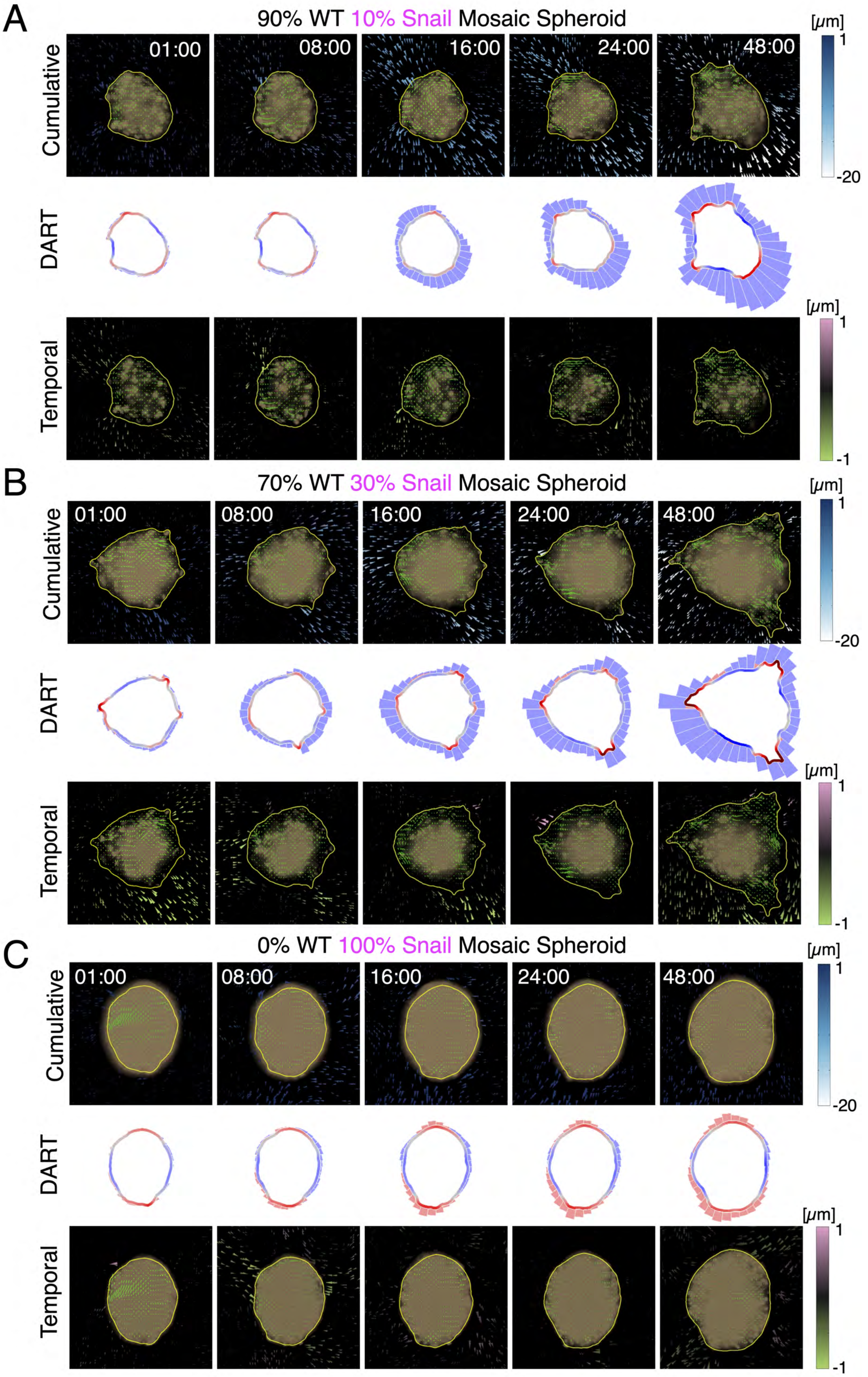




FIG. S10. **(A)** Cumulative, DART and temporal particle displacements for 90% wildtype and 10% Snail mosaic spheroid. **(B)** Cumulative, DART and temporal particle displacements for 70% wildtype and 30% Snail mosaic spheroid. **(C)** Cumulative, DART and temporal particle displacements for 100% Snail spheroid.

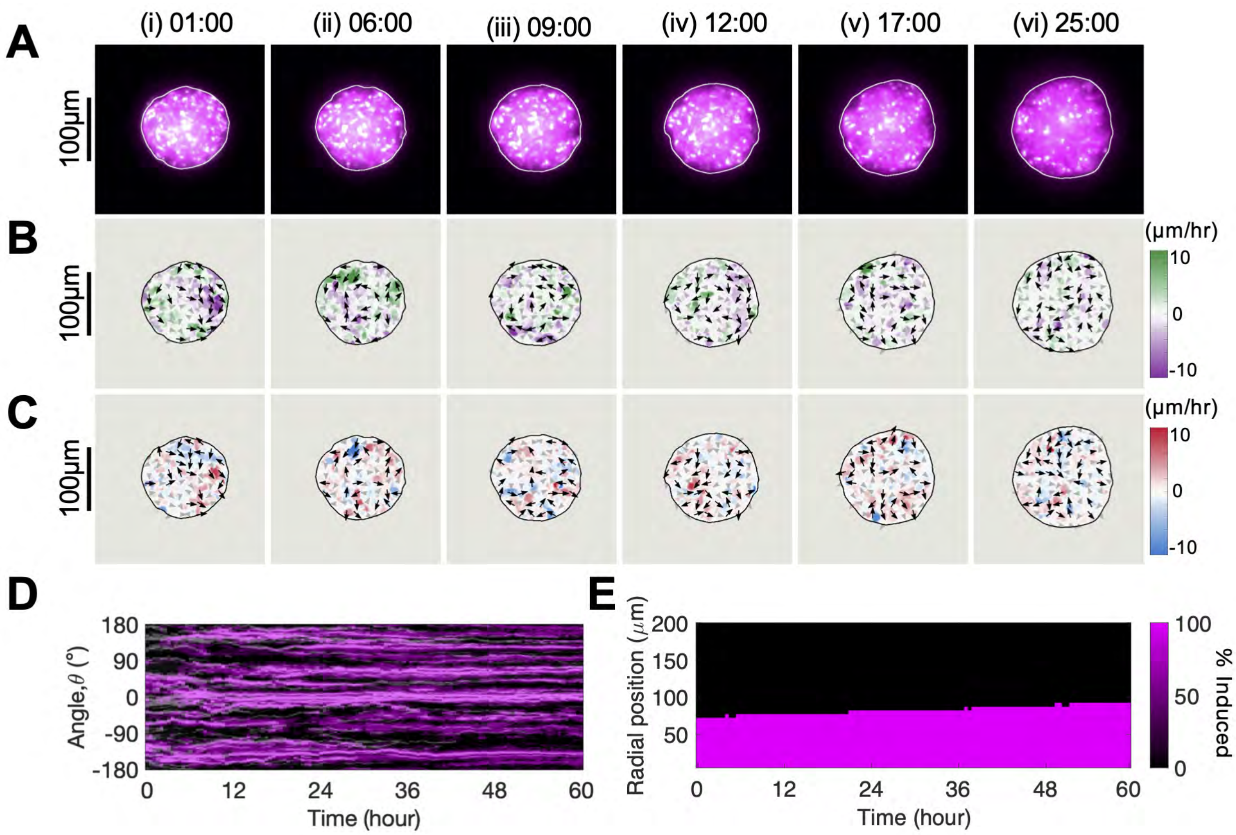


FIG. S11. **Minimal invasion by 100% Snail spheroids**. **(A)**Representative snapshots of 100% Snail spheroids. **(B, C)** spheroids exhibit lower and uncoordinated circumferential and radial velocities. **(D)** Kymograph shows limited and uncoordinated angular motion of Snail cells in the outermost layer from 0-60 h. **(E)** Snail cells are uniformly distributed throughout the spheroid.

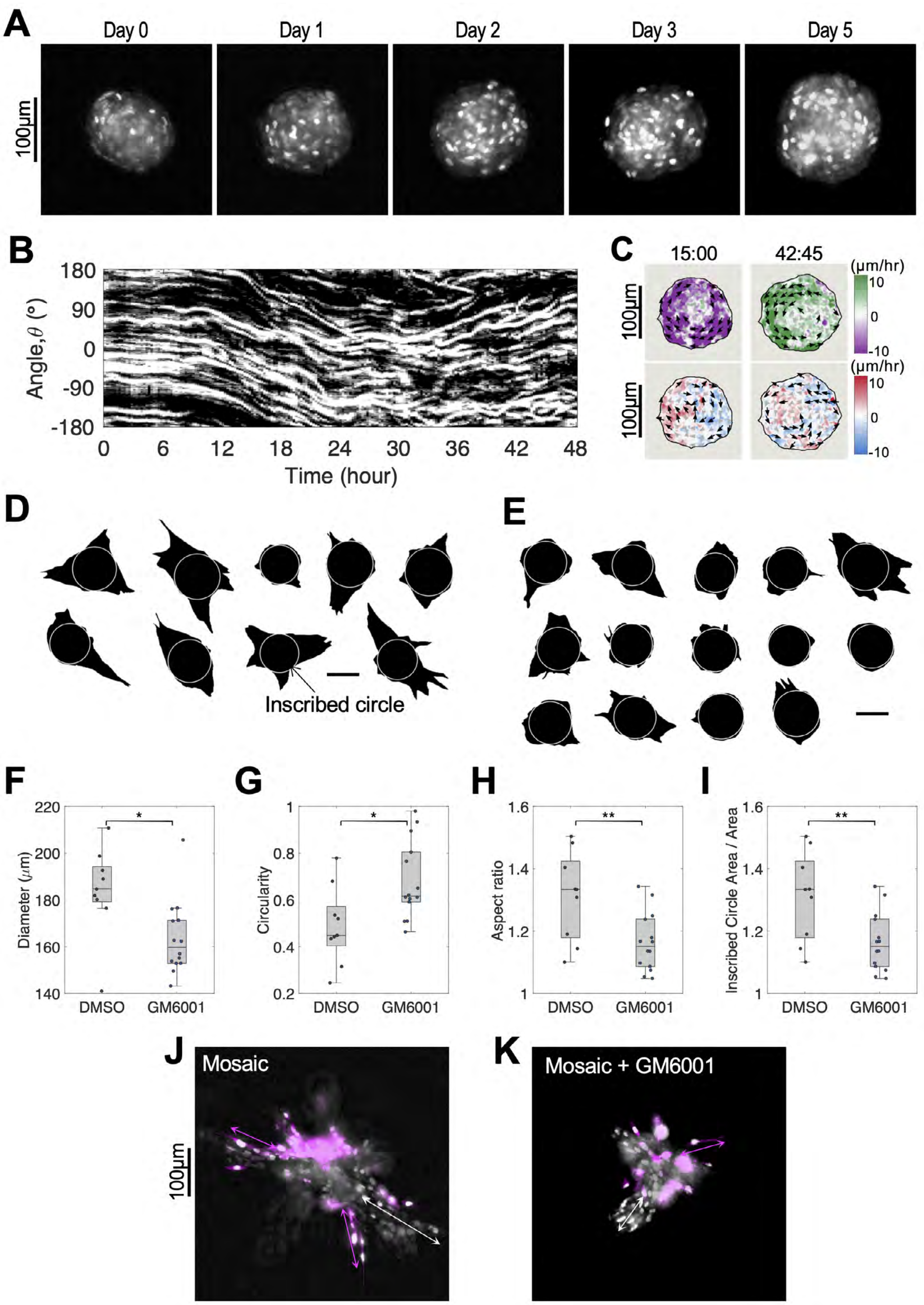


FIG. S12. **MMP inhibitor GM6001 maintains orbiting and limits invasion**. **(A)** Representative spheroid morphologies over time under 25 $\mu$M GM6001 condition. **(B)** Kymograph of the outermost layer of GM6001-treated spheroid, showing a switch in orbiting direction after approximately 24 h. **(C)** Representative velocity profiles of the GM6001-treated spheroid. The spheroid exhibited coordinated orbiting in clockwise direction at 15 h but switched to counterclockwise at 42.75 h. No significant outward radial velocity hotspot was observed at either time point. **(D, E)** Representative images of DMSO control and GM6001-treated spheroids at day 5, respectively. White circles indicate the inscribed circles. **(F-I)** Quantification of diameter, circularity, aspect ratio, and inscribed circlearea of DMSO control and GM6001 spheroids. GM6001 spheroids were significantly smaller and rounder. The larger portions of inscribed circle area relative to the spheroid area indicates fewer and less invasive protrusions from the treated spheroids. **(J,K)** 30% Mosaic spheroids under DMSO control, and GM600l treated conditions. Mosaic spheroid under control condition developed signficant invasive protrusions whereas the GM6001 spheroids exhibited limited invasion, confirming that the MMP inhibition affect not only WT cell collective migration but also the Snail-cell-driven collective migration. Two-tailed t-test, $*p < 0.05$.

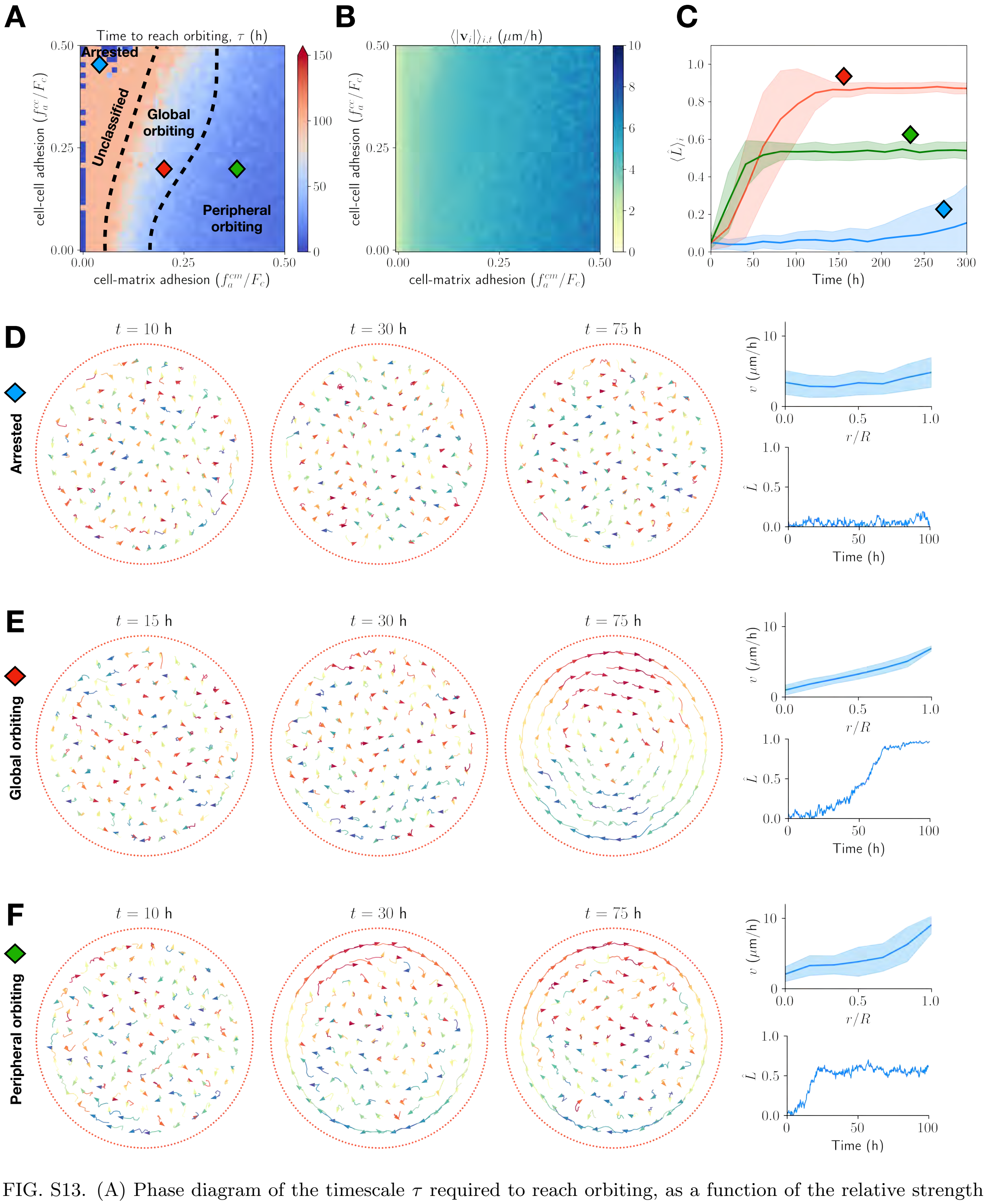


FIG. S13. (A) Phase diagram of the timescale $\tau$ required to reach orbiting, as a function of the relative strength of cell-cell and cell-matrix adhesion. Timescales are estimated by fitting a sigmoid-like function to the time-dependent average normalized angular momentum $\langle\hat{L}\rangle_i$ (see SI for details). (B) Phase diagram of mean cell speed at equilibrium, which depends on active-drag and cell-matrix forces. For fixed self-propulsion ($\alpha$) and friction ($\beta$) parameters, cell speed increases with cell-matrix adhesion. (C) Angular momentum order parameter dynamics in three distinct regions of the phase diagram: arrested motion (blue), global orbiting (red), and peripheral orbiting (green). (D-F) Representative snapshots of the three behaviors predicted by the model (arrested motion, global orbiting, and peripheral orbiting) at $t$ = 10, 30, 75 h, along with mean cell speed as a function of spheroid radius and angular momentum order parameter dynamics.

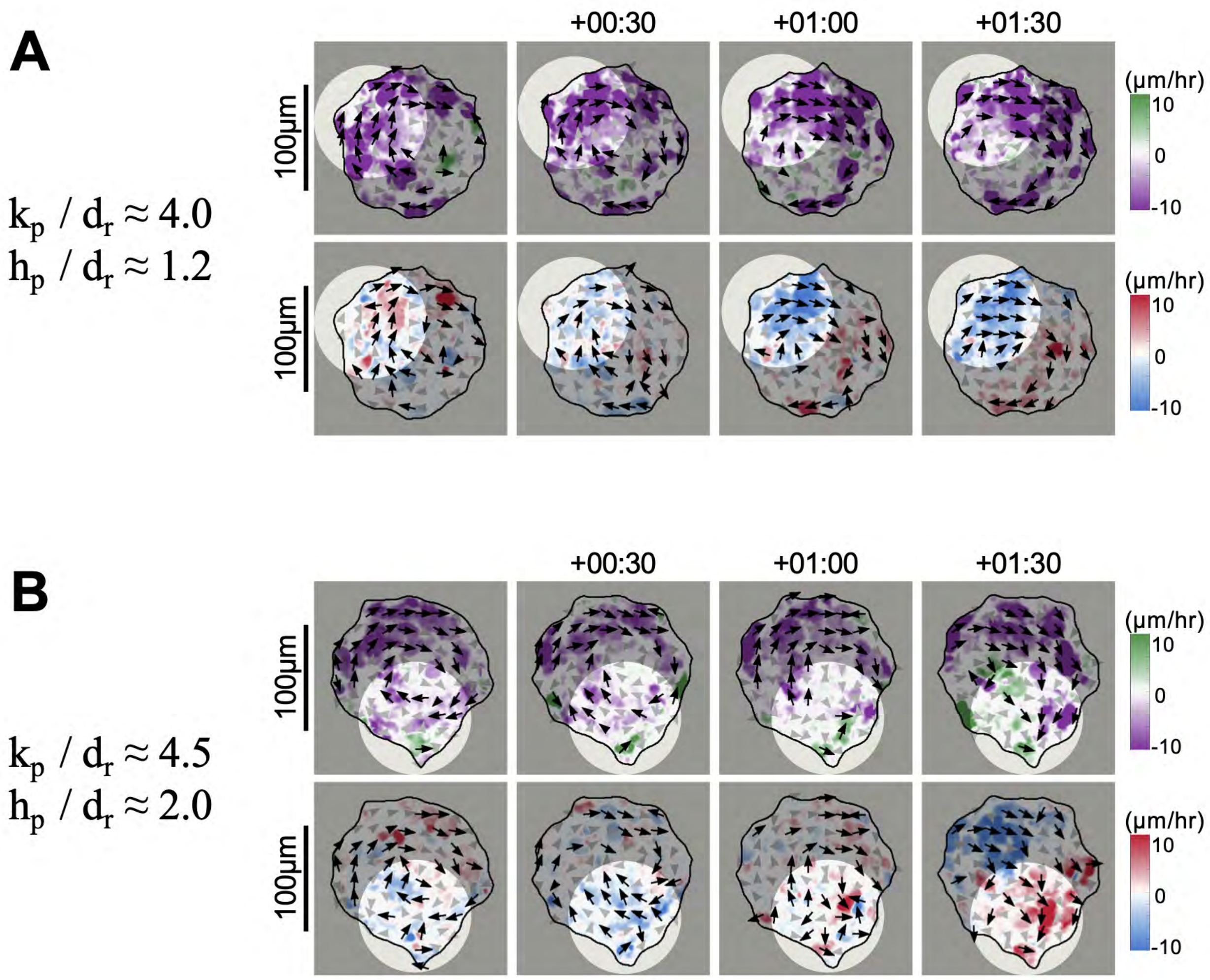


FIG. S14. **Geometry of bump perturbation governs orbiting persistence**. **(A)** Velocity profiles from experimental data of spheroids with a comparable perturbation size of $k_p/d_r = 4$ and $h_p/d_r = 1.2$. The orbiting cells were able to overcome the small perturbation and maintain orbiting. **(B)** The orbiting motion of spheroid with a larger perturbation, $k_p/d_r = 4.5$, and $h_p/d_r = 2$ was impeded by the geometrical effect. Cells near the perturbation exhibit limited migratory speed (arrested) and eventually begin to migrate outward.

### Appendix A: Mathematical Model

#### 1. Model Description

To gain better intuition into how forces between cells and their ECM boundary affect collective orbiting, we developed a computational model with tunable mechanistic interactions and analyzed how perturbations to this system affected the stability of coordinated rotation. In this minimal 2D model, cells are treated as active, self-propelled agents that interact with each other via soft repulsion at short distances, which ensures that individuals cannot occupy the same spatial location, and adhesion-based attraction mediated by cell protrusions at longer distances [8]. It was previously shown in this model that collective orbiting ("milling") arises among agents in the absence of boundaries when their cell-cell forces are defined by certain potentials [9].

We treat cells as discrete particles moving off-lattice in a 2D domain. Our decision to use a 2D model is motivated by the observation that cell movement between different layers of the spheroids appears to be limited, however we note that the framework described below can be extended naturally to 3D. Cells are treated as active particles subject to drag (frictional) forces. We incorporate two key terms to describe these dynamics: a self-propulsion term proportional to the velocity of cell $i$, and a nonlinear drag (friction) term which acts against the cell velocity and is proportional in magnitude to the cube of the cell speed. This formulation, commonly used in models of self-propelled particles as a first approximation for active movement, naturally predicts an equilibrium speed at which cells travel where friction balances self-propulsion such that the speed of cell $i$ is given by $|\mathbf{v}_i| = \sqrt{\alpha/\beta} =: u$ in the absence of other forces.

We assume that cells can interact with each other and with their ECM boundary via cell-cell, $\mathbf{F}^{cm}$, and cell-matrix, $\mathbf{F}^{cm}$, forces which each encode soft repulsion at short distances and adhesion-based attraction at longer distances. The strength and magnitude of the cell-cell (*resp.* cell-matrix) force is assumed to depend on the relative displacement between the cell and its neighbor (*resp.* neighboring ECM molecule). The cell-matrix force additionally depends on the direction in which the cell travels, as we will later introduce the assumption that cells can only adhere to ECM molecules within a specified angle of this direction. The specific functions that yield the cell-cell and cell-ECM forces are described in more detail in the ensuing paragraphs.

We model the ECM boundary as a set of discrete equally spaced particles to simplify numerical calculations of cell-matrix forces and facilitate the introduction of boundary perturbations. This modeling approach further admits straightforward interpretation of how specific ECM molecules affect cell movement. For a circular boundary of radius $R$ made up of $M$ points, the position of the $l^{\text{th}}$ ECM molecule is given by $\mathbf{y}_l = (R\cos 2\pi l/M, R\sin(2\pi l/M))$ for $l = 1, \ldots, M$. In the main text of this manuscript, we assume that these points are static so that we may focus on how the boundary geometry, cell-cell, and cell-matrix force influence the stability of collective orbiting in a controlled setting.

We use Newton's second law to supply ordinary differential equations (ODEs) dictating how the position and velocity of cell $i$, $\mathbf{x}_i(t)$ and $\mathbf{v}_i$, respectively, change over time $t \geq 0$ in response to the active, drag, cell-cell, and cell-matrix forces. The system of ODEs is given by

$$\frac{\mathrm{d}\mathbf{x}_i}{\mathrm{d}t} = \mathbf{v}_i\,, \tag{A1}$$

$$\frac{\mathrm{d}\mathbf{v}_i}{\mathrm{d}t} = \underbrace{\left(\alpha - \beta|\mathbf{v}_i|^2\right)\mathbf{v}_i}_{\text{active and drag forces}} + \underbrace{\sum_{j\neq i,\, j=1}^{N} \mathbf{F}^{cc}(\mathbf{x}_j - \mathbf{x}_i)}_{\text{cell-cell repulsion/attraction}} + \underbrace{\sum_{k=1}^{M} \mathbf{F}^{cm}(\mathbf{y}_k - \mathbf{x}_i, \mathbf{v}_i)}_{\text{cell-ECM repulsion/attraction}}\quad, \tag{A2}$$

$$\frac{\mathrm{d}\mathbf{y}_k}{\mathrm{d}t} = \mathbf{0}\,, \tag{A3}$$

---

[8] D'Orsogna, M.R., et al. *Phys Rev Lett.* 96(10), 104302 (2006); Bertozzi, A.L., et al. *Commun Math Sci.* 13(4), 955-985 (2015); Carrillo, JA, et al. *J. Theor. Biol.* 445, 75-91 (2018)

[9] Albi, G., et al. *SIAM J. Appl. Math.* 74(3), 794-818 (2014).

where $\mathbf{y}_l$ denotes the position of the $l^{\text{th}}$ ECM molecule. We have slightly abused notation by absorbing the mass of the cell into the terms on the right-hand side of Eqs. (A1)-(A2).

We consider the direction of cell-cell and cell-matrix forces to be aligned with the displacement vector, $\mathbf{r}$, between a cell and its neighboring cell or ECM molecule. The magnitudes of both forces are further assumed to only depend on the distance between these two objects. This is captured by using two radially symmetric kernels, $K^{cc}(|\mathbf{r}|)$ and $K^{cm}(|\mathbf{r}|)$, which are related to the forces via

$$\mathbf{F}^{cc}(\mathbf{r}) = K^{cc}(|\mathbf{r}|)\frac{\mathbf{r}}{|\mathbf{r}|}, \tag{A4}$$

$$\mathbf{F}^{cm}(\mathbf{r},\mathbf{v}) = \tilde{H}(|\mathbf{r}|,\, \hat{\mathbf{r}}\cdot\hat{\mathbf{v}} - \cos\theta_{\max})\, K^{cm}(|\mathbf{r}|)\frac{\mathbf{r}}{|\mathbf{r}|}\,. \tag{A5}$$

The term which is unique to the cell-ECM force, $\tilde{H}(\cdot,\cdot)$, is a modified Heaviside function that arises from an assumption that cells can only adhere to matrix molecules that fall within an angle $\theta_{\max} \in [0,\pi]$ of their unit velocity vector $\hat{\mathbf{v}}$. Repulsive forces are not assumed to be affected by the modified Heaviside function, as they are assumed steric in origin and therefore isotropic. The assumption of anisotropic cell-ECM adhesion accelerates the emergence of collective orbiting, but we show in Fig. S18 that orbiting robust across a large array of values for $\theta_{\max}$. The modified Heaviside function is defined as

$$\tilde{H}(r,\,\lambda) = \begin{cases} 1, & \text{if } 0 < r < d_a; \\ H(\lambda), & \text{if } r \geq d_a\,; \end{cases}$$

where $H(\cdot)$ is the standard Heaviside function.

In all numerical simulations, we fix the angular sensing region such that the cells do not adhere to ECM molecules which lie outside an angle $\theta_{\max} = \pi/3$ from its current velocity vector. In Fig. S18 we quantify the impact of this parameter, showing that results are robust under similar values. Note also that cell-matrix adhesion changes the equilibrium velocity of cells, such that the long-term cell speed will be slightly larger than $u = \alpha/\beta$.

For simplicity we assume that repulsion ($f_r$) is identical for cell-cell and cell-matrix interactions, while adhesion depends on two parameters, $f_a^{cc}$ and $f_a^{cm}$, that correspond to the respective kernel. While there are multiple possibilities for the functional forms of the repulsive and attractive kernels, for example power-law or exponentially decaying functions [10], we opt instead to use kernels which ensure that cells do not experience forces from objects which lie beyond a certain maximum distance. The cell-cell and cell-matrix kernels used in this manuscript are based on [39], which have compact support and have been well characterized. They are given by

$$K^{\ell}(r) = \begin{cases} -f_r\left(\dfrac{d_r}{2r}\right)^{3-2s}, & \text{if } 0 < r < \frac{d_r}{2}; \\ \dfrac{2f_r(r-d_r)}{d_r}, & \text{if } \frac{d_r}{2} \leq r < d_r; \\ -\dfrac{4f_a^{\ell}(r-d_a)(r-d_r)}{(d_a-d_r)^2}, & \text{if } d_r \leq r < d_a; \\ 0, & \text{if } r \geq d_a, \end{cases} \tag{A6}$$

where $\ell \in \{cc, cm\}$, $s \in (0,2]$ is a parameter that captures the behavior of the kernel near the origin and is related to the cell nucleus stiffness, $d_r$ is a typical cell diameter, and $d_a$ gives the range of adhesive interactions. For the simulations in the main text, we set $s = 1.25$, which corresponds to a singular kernel at the origin. In the sections below, we show that a different choice of the exponent $s$ affects the results only qualitatively. This is consistent with previous work using the same family of kernels [11]. Fig. S15 shows a plot of this kernel.

---

[10] Albi, G., et al. *SIAM J. Appl. Math.* 74(3), 794-818 (2014); D'Orsogna, M.R., et al. *Phys Rev Lett.* 96(10), 104302 (2006
[11] Carrillo, JA, et al. *J. Theor. Biol.* 445, 75-91 (2018)

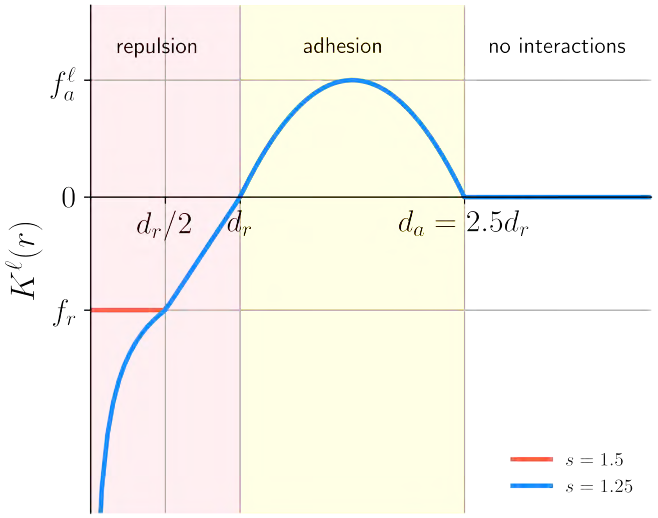


FIG. S15. **Representation of the interaction kernels $K^\ell(r)$.** Parameters are not to scale.

#### *a. H-stability and adhesion parameter ranges*

We use the concept of H-stability from statistical mechanics to constrain the range of admissible model parameters [39]. This constraint ensures that cells do not "collapse" to a single point as more individuals are added to the simulation. In biological terms, this means that there is a sufficient balance between repulsive and attractive forces such that, as more individuals are added into a simulation space, cells do not form clusters with a distance less than a cell diameter. If the kernels are not H-stable, then unphysical solutions can arise in which a very large number of particles occupies a small, finite region of space.

Mathematically, a kernel or potential is H-stable if the total potential energy of the system is bounded below by a constant that is independent of the number of particles. This concept has been applied in the context of interacting models of cell migration and is well-characterized for the kernel in Eq. (A6) [12]. (This is another motivating reason for why we have used the kernels described abov). In particular, given an interaction potential $u_\ell(r)$, such that $K^\ell(r) = u_\ell'(r)$, $K^\ell$ is called H-stable if

$$\int_0^{+\infty} u_\ell(r) r \, \mathrm{d}r > 0 \, .$$

If the integral is negative, then the kernel is not H-stable (also called "catastrophic"). For the kernel in Eq. (A6), the integral above yields the following constraint on the strength of repulsive and attractive forces:

$$\frac{f_r}{f_a^\ell} > \frac{32s(d_a - d_r)(3d_a^2 + 4d_a d_r + 3d_r^2)}{5(11s+6)d_r^3} := F_* \, . \tag{A7}$$

Following these ideas, in numerical simulations we vary the cell-cell and cell-matrix adhesion strengths in $f_a^\ell \in [0, F_c]$, with $F_c := f_r/F_*$ so that the potential is always H-stable.

To illustrate the impact of H-stability, we present snapshots of numerical simulations in which the kernels are not H-stable — see Fig. S16. Depending on the H-stability of the cell-cell and cell-matrix interactions, the system either collapses to the center of the spheroid or to its boundary. Taken together with the above simulations, these results indicate that a balance of cell-cell and cell-matrix forces is necessary to stabilize a collective orbiting state.

---

[12] Carrillo, JA, et al. *J. Theor. Biol.* 445, 75-91 (2018)

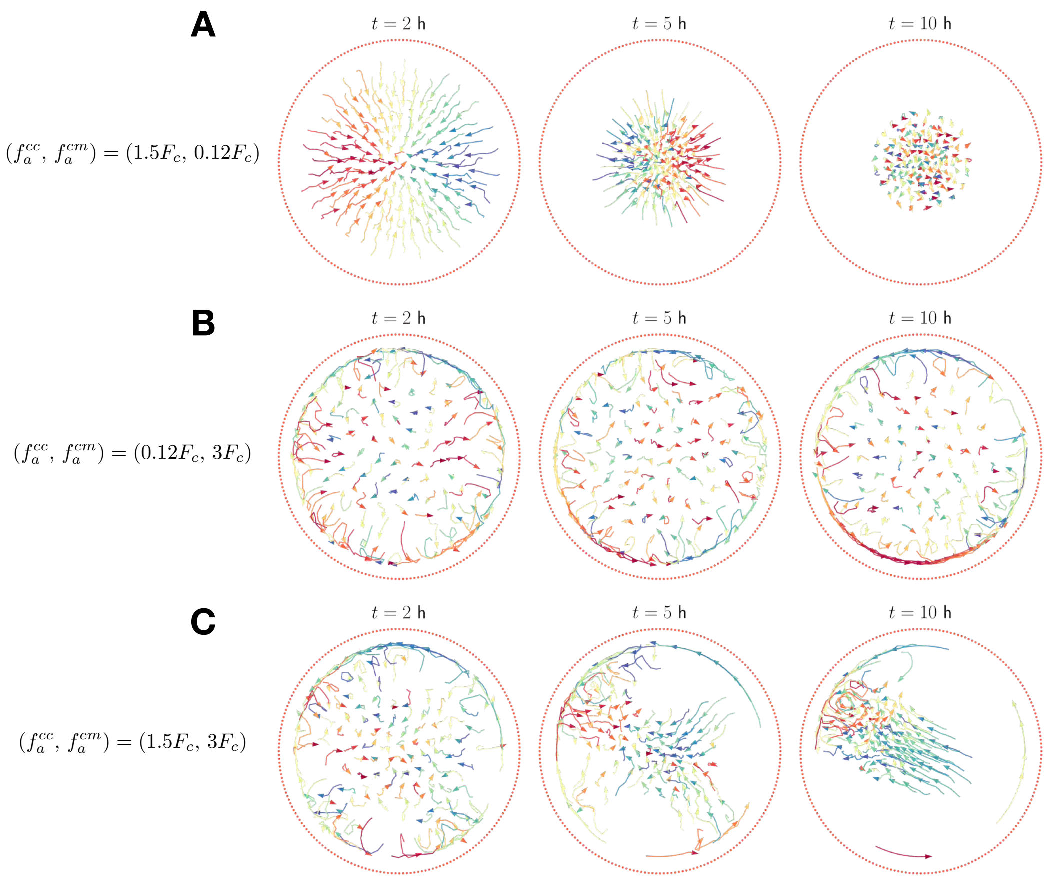


FIG. S16. **Interaction kernels that are not H-stable lead to cell collapse and unphysical behavior.** Snapshots of numerical simulations where: (A) $K^{cc}$ is not H-stable; (B) $K^{cm}$ is not H-stable; and (C) neither is H-stable.

#### *b. Model parameters*

We set the radius of the circular ECM boundary, $R = 100$ $\mu$m, to have a comparable length scale to experimentally observed spheroid radii. For simplicity, the length scales of repulsive and adhesive forces, $d_r$ and $d_a$, and the strength of repulsive forces $f_r$, are assumed to be identical for cell-cell and cell-matrix interactions. The parameter $d_r$ can be estimated from cell-cell spacing observed in experiments, while $d_a$ can be similarly estimated by noting that MCF-10A cells do not appear to interact with neighbors beyond $2-3$ cell diameters. This leads us to fix $d_r = 15$ $\mu$m, $d_a = 2.5d_r = 37.5$ $\mu$m. The experimental data also provide sufficient information to fix the asymptotic cell speed, which is roughly equal to $u = \sqrt{\alpha/\beta} = 3$ $\mu$m/h. The average cell velocity in numerical simulations, however, is higher than $u$ —around 10 $\mu$m/h— due to the additional tangential acceleration provided by cell-matrix adhesion. Furthermore, we note that fixing the cell speed determines the ratio of the parameters $\alpha$ and $\beta$ but not their specific values.

The number of discrete ECM particles, $M$, is chosen to be sufficiently large to avoid cell escape from the spheroid, yet also small enough to ensure that the net cell-matrix force on a given cell is no more than an order of magnitude larger than the cell-cell repulsion force. We have found that, for the parameters used in this manuscript, $M = 200$ molecules is sufficient to achieve both goals. For a circular ECM boundary of radius $R = 100$ $\mu$m, this corresponds to a one-dimensional ECM density of 0.32 molecules/$\mu$m, which is approximately five times larger than the one-dimensional density corresponding to the external layer of orbiting cells.

TABLE S1. **Summary of model and simulation parameters and their values used in numerical simulations.**

| Parameter | Values | Units | Meaning |
|---|---|---|---|
| $\alpha$ | $10^{-2}$, $10^{-1}$ (main text), $10^{0}$ | $\mathrm{h}^{-1}$ | self-propulsion coefficient |
| $u$ | 3 | $\mu$m/h | equilibrium cell speed in the absence of cell-cell and cell-matrix forces |
| $\beta$ | $\alpha/u^2$ | h/$\mu$m$^2$ | friction coefficient |
| $d_r$ | 15 | $\mu$m | repulsion interaction range |
| $d_a$ | $2.5d_r$ | $\mu$m | adhesion interaction range |
| $f_r$ | 100 (main text) | $\mu$m/h$^2$ | cell-cell and cell-matrix repulsion strength |
| $F_c$ | 5.18 (main text) | $\mu$m/h$^2$ | critical adhesion strength for H-stability (Eq. (A7)) |
| $f_a^{cc}$ | $[0, 0.5F_c]$ (main text), $1.5F_c$ (Fig. S16) | $\mu$m/h$^2$ | cell-cell adhesion strength |
| $f_a^{cm}$ | $[0, 0.5F_c]$ (main text), $3F_c$ (Fig. S16) | $\mu$m/h$^2$ | cell-matrix adhesion strength |
| $\theta_{\max}$ | $\pi/3$ (main text), $[0, \pi]$ (Fig. S18) | rad | maximum cell-matrix alignment angle (Eq. (A5)) |
| $s$ | 1.25 | — | exponent controlling the repulsive force behavior at the origin |
| $\rho$ | 0.004 | cells/$\mu$m$^2$ | cell density |
| $N$ | 130 (circular boundary) $+ n_p\rho\Delta A$ (boundary perturbations, Eq. (A12)) | cells | number of cells |
| $M$ | 200 | molecules | number of ECM molecules |
| $R$ | 100 | $\mu$m | spheroid radius |
| $h_p$ | $[0, 40]$ | $\mu$m | height of boundary perturbation |
| $k_p$ | $[0, 80]$ | $\mu$m | width of boundary perturbation |
| $n_p$ | $1, 2, 3$ | — | number of boundary perturbations |

The number of cells, $N$, is similarly chosen to maximize the number of individuals in the spheroid while simultaneously limiting the degree of overlap between them, as otherwise repulsive forces overly dominate cell movement and uncoordinated behavior occurs. Since cells are assumed to be spherically symmetric, the problem reduces to finding the maximum number of identical circles that fit within a unit circle without overlapping. Although this classical problem only has explicit solutions for relatively small cell numbers [13], we note that the density of circles appears to asymptotically approach a constant value of $0.75 - 0.85$ as the number of cells increases. For circles of radius $d_r/2 = 7.5$ $\mu$m embedded in a larger circle of radius $R = 100$ $\mu$m, this density is approximately equivalent to $130 - 140$ cells. Consequently, we set $N = 130$ cells for the simulations.

We have thus far fixed all but four parameters in the mathematical model, with $\alpha$, $f_r$, $f_a^{cc}$, and $f_a^{cm}$ remaining undetermined. The adhesion-related parameter values are constrained by Eqn. (A7), ensuring that the cell-cell and cell-matrix potentials are H-stable. As it is unclear what are reasonable values for these parameters, we sweep over various values in the main text to determine how they influence the long-term system behavior. Meanwhile, the parameter $f_r$ sets the timescale to reach the equilibrium orbiting scale. All simulations in the main text have $f_r = 100$ $\mu$m/h$^2$ as we observe in numerical simulations that this corresponds to an equilibrium timescale of around $10 - 20$ h. Finally, $\alpha^{-1}$ represents the timescale at which cells readjust their velocities as a consequence of active and drag forces. To prevent cells from escaping the spheroid, the acceleration caused by cell-matrix repulsion must dominate over the acceleration arising from active-drag forces, which drives changes in the velocity $\Delta v \sim u$ during timescales approximately equal to $\alpha^{-1}$. Consequently, the condition $f_r \gg u\alpha$ is fulfilled, and we set $\alpha = 0.01, 0.1, 1\,\mathrm{h}^{-1}$, all of which satisfy this requirement.

We summarize in Table S1 the model parameters and their chosen values. We also study in the next sections the impact of varying different model parameters.

---

[13] Graham, R.L., et al. *Discrete Mathematics.* 181(1-3), 139-154 (1998).

#### *c. Numerical implementation*

We solve the system of ODEs given by Eqns. (A1)-(A3) in Python, using an explicit fourth order Runge-Kutta scheme with time step $\Delta t = 0.1$h. Initial cell positions were sampled uniformly at random within a circle of radius $R = 100$ $\mu$m and with a minimum interparticle distance of 10 $\mu$m. Initial velocities were similarly sampled uniformly at random, with each component of the velocity sampled uniformly on the interval $[-\sqrt{10}, \sqrt{10}]$ $\mu$m/h.

### 2. Angular momentum

We quantify orbiting with using the normalized average angular momentum, $\hat{L}$, as an order parameter. Its value is defined as

$$\hat{L} = \frac{1}{N}\left|\sum_{i=1}^{N} \hat{\mathbf{x}}_i \times \hat{\mathbf{v}}_i\right| , \tag{A8}$$

where $\hat{\mathbf{x}}_i$ denotes the unit position vector of cell $i$ and $\hat{\mathbf{v}}_i$ its unit velocity vector. Note that, with this definition, we have assumed the system center of mass is located at the origin. This order parameter satisfies $\hat{L} \geq 0$ and also

$$\hat{L} \leq \frac{1}{N}\sum_{i=0}^{N} |\hat{\mathbf{x}}_i \times \hat{\mathbf{v}}_i| = 1 .$$

with equality when each position vector is orthogonal to its corresponding velocity vector. Notably, this occurs in a perfectly coordinated state where all cells orbit either clockwise or anticlockwise. Hence, values of the angular momentum close to unity indicate global orbiting. By contrast, order parameter values close to zero are indicative that most cells do not exhibit collective coordination.

#### *a. Phase diagrams*

To produce the phase diagrams in the main text, we vary the strengths of cell-cell and cell-matrix adhesion, $f_a^{cc}$ and $f_a^{cm}$, respectively, within the interval $[0, 0.5F_c]$, as predicted by the H-stability analysis. These values are chosen because, for strong cell-cell adhesion ($f_a^{cc} > 0.5F_c$), we observed parameter combinations that do not provide sufficient coverage of the spheroid. The space $[0, 0.5F_c] \times [0, 0.5F_c]$ is discretized into a $40 \times 40$ grid, and fifty simulations corresponding to 300 h are performed for each parameter combination. All other parameters are kept fixed, and numerical simulations differ only in their initial conditions. We then average the normalized angular momentum over the last 100 h of the numerical simulation and across the fifty different simulations. We denote by $\langle\cdot\rangle_i$ averages with respect to the simulation replicates, and by $\langle\cdot\rangle_t$ the averages with respect to time.

Numerically, we observe that cell-matrix interactions drive the emergence of collective orbiting within the first few hours of the simulation. In contrast, when cell-matrix adhesion is weak, collective orbiting does not appear within the first $\sim 100-200$ h. However, in a small fraction of simulations, global orbiting eventually emerges over much longer timescales. In these cases, the average normalized angular momentum, $\langle\hat{L}\rangle_{i,t}$, takes up to $\sim 10^3$ h to reach equilibrium. Here, collective orbiting arises purely due to geometric boundary constraints, as cell-matrix adhesion is almost negligible, leading to a significantly longer coordination timescale. To identify such cases, we check whether the increase in normalized angular momentum during the last 100 h is greater than 0.1 — corresponding to a slope of more than 0.001 in a linear regression fit. Simulations meeting this criterion are labeled as *unclassified* and appear white in the phase diagram.

Overall, we found that an uncoordinated migration regime occurs in the limit of weak cell-matrix adhesion and strong cell-cell adhesion (Fig. 6AB), as the angular order parameter is well below 0.2 and the average root mean squared cell speed is approximately 2 $\mu$m/h, well below the equilibrium cell speed of 8-10 um/h

(Fig. S12). Globally coordinated orbiting occurs in an intermediate regime with slightly weaker cell-matrix adhesion and slightly stronger cell-cell adhesion, where $\langle L \rangle \sim 0.8 - 1.0$. Analysis of the spatial distribution of cell speeds for this regime indicate that individuals at the periphery of the spheroid migrate faster on average compared to agents within the interior (Fig. S12). Further increases in the cell-matrix adhesion strength parameter result in a decrease of the angular order parameter, with $\langle L \rangle \sim 0.4 - 0.6$. Subsequent investigation of individual realizations reveals that this decrease is due to uncoordinated migration of cells within the spheroid interior, whereas agents located at the periphery maintain circumferential orbiting (Fig. 6AB).

#### b. *Timescale to reach orbiting*

To quantify the time to reach the orbiting state, $\tau$, we first average $\hat{L}$ across the simulation replicates to obtain $\langle \hat{L} \rangle_i$. We then fit $\langle L \rangle_i$ as a function of time, to a sigmoid function using non-linear least squares. The sigmoid function, defined as

$$\hat{L}(t) = L_0 + \frac{L_{\text{eq}}}{1 + \exp\left(-(t-\tau)/b\right)}, \tag{A9}$$

depends on time, $t$, and on four parameters: the initial angular momentum, $L_0$; the equilibrium angular momentum, $L_{\text{eq}}$; the time at which $\hat{L} = L_0 + L_{\text{eq}}/2$, $\tau$; and a parameter that quantifies the steepness of the sigmoid, $b$. The fitted parameter ($\tau$) is shown in Fig. S19, along with representative snapshots of numerical simulations and their corresponding average angular momentum dynamics.

#### c. *Scaling with number of orbiting layers*

In experiments and numerical simulations we often observe states in which only cells within the external layers of the spheroid orbit, while cells near the center remain uncoordinated. To quantify this, we study how the angular momentum scales with the number or orbiting layers, $n_{\text{orb}}$. Given the typical distance between cells, $a$, and the outer radius of the spheroid, $R$, $n_{\text{orb}}$ is a non-negative integer between 0 and $R/a$. Next, we observe that the number of orbiting cells in a layer of radius $r_j$ is approximately $2\pi r_j/a$, where the allowed values for $r_j$ are given by $r_j = R - ja$, with $j = 1, \ldots, n_{\text{orb}}$. By assuming that uncoordinated cells negligibly influence the total angular momentum of the system and all cells within orbiting layers are perfectly aligned, we can approximate Eq. (A8) as

$$\hat{L} \approx \frac{2\pi}{N} \frac{(R-a) + (R-2a) + \ldots + (R - n_{\text{orb}}a)}{a} = \frac{2\pi n_{\text{orb}}}{N}\left(\frac{R}{a} - \frac{n_{\text{orb}}+1}{2}\right). \tag{A10}$$

When the maximum number of possible layers $n_{\text{orb}} = R/a \gg 1$, this approximation further reduces to

$$\hat{L} \approx \frac{2\pi R}{Na}\left(\frac{R}{2a} - \frac{1}{2}\right) \approx \frac{\pi R^2}{Na^2} = 1\,,$$

so it is consistent with the expected angular momentum value for a global orbiting state.

Identifying the typical distance between cells as $a \sim \rho^{-1/2} = R\sqrt{\pi/N}$, Eq. (A10) becomes independent of the geometry of the spheroid

$$\hat{L} \approx \frac{2\pi n_{\text{orb}}}{N}\left(\sqrt{\frac{N}{\pi}} - \frac{n_{\text{orb}}+1}{2}\right). \tag{A11}$$

In particular, for our numerical simulations with $N = 130$, the normalized angular momentum should correspond to $\hat{L} \sim 0.26, 0.48, 0.64, 0.76$ for one, two, three, and four layers, respectively. We use this criterion to generate a discretized version of the phase diagram in the main text (Fig. 6B), classifying states into four categories based on whether $\langle L \rangle_i$ reaches equilibrium within 200 h and the approximate number of orbiting layers (see Fig. S17). States labeled as arrested do not exhibit orbiting within 300 h of simulation but may transition to an orbiting state over much longer timescales.

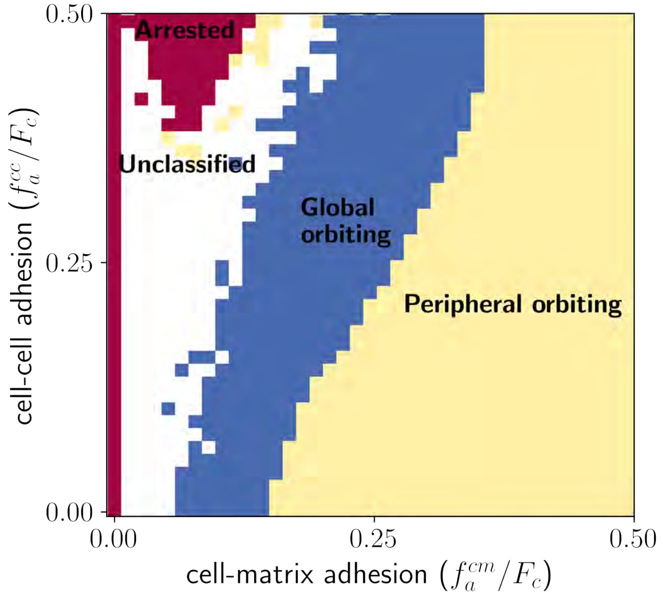


FIG. S17. **The mathematical model predicts three regimes based on collective orbiting behavior.** States are classified according to the number of orbiting layers, as determined by the normalized angular momentum: arrested ($\langle\hat{L}\rangle_{i,t} < 0.24$, no layers), global orbiting ($0.24 \leq \langle\hat{L}\rangle_{i,t} < 0.76$, one to three layers), and peripheral orbiting ($\langle\hat{L}\rangle_{i,t} \geq 0.76$, four or more layers). Unclassified states reach equilibrium only after very long times.

#### 3. Boundary perturbations

We then considered the stability of collective orbiting to outward-oriented Gaussian perturbations of the ECM boundary with height $h_p$ and width $k_p$. We found that collective orbiting remained relatively stable when the boundary was perturbed by only one sharp feature, with cells particularly unaffected when the perturbation height was less than 1 cell diameter and its width less than 2 cell diameters (Fig. 6CD). Upon observing individual snapshots of these simulations, it is apparent that most cells collectively orbit by following the nearly constant curvature of the boundary and are only slightly disrupted by misaligned cells near the perturbation. By contrast, two or more sharp features were sufficient to destabilize collective orbiting when their height exceeded one cell diameter and width exceeded two cell diameters (Fig. 6EF and Fig. S20). The presence of two or more sharp perturbations both presents increased obstacles to coordinated migration and decreases the length of continuously connected curved boundary. Thus, mechanistic cell-matrix interactions and the geometry of the boundary interface are sufficient to explain the experimental observation that radial invasion in spheroids is typically associated with at least two strands, as otherwise the collective orbiting state is preferred.

All numerical simulations in the previous sections used a circular matrix boundary, given by the position of discrete particles $\mathbf{y}_l = (R\cos 2\pi l/M, R\sin(2\pi l/M))$ for $l = 1, \ldots, M$. Here, we introduce boundary perturbations with height $h_p$, and width $k_p$. The perturbed circle can be parametrized by $\theta \in [0, 2\pi]$, with the peak of the perturbation found at $\theta = \pi$:

$$\mathbf{y}(\theta) = R_p(\theta)\,(\cos\theta,\, \sin\theta)\,, \quad R_p(\theta) = R + h_p \exp\{-4R^2(\theta-\pi)^2/k_p^2\}\,.$$

The case of boundary perturbations at $n_p$ different locations can be considered analogously. In numerical simulations, we reparametrize the perturbed boundary by the arc length parameter to generate matrix particles with a fixed density ($200/2\pi R \approx 0.32$ cells/$\mu$m).

#### a. Change of area due to perturbation

To keep constant cell density inside the spheroid, we calculate the change in area due to one boundary perturbation

$$\begin{aligned}\Delta A &= \frac{1}{2}\int_0^{2\pi} R_p(\theta)^2\,\mathrm{d}\theta - \pi R^2 \\ &= R r_p \int_0^{2\pi} e^{-4R^2(\theta-\pi)^2/k_p^2}\,\mathrm{d}\theta + \frac{r_p^2}{2}\int_0^{2\pi} e^{-8R^2(\theta-\pi)^2/k_p^2}\,\mathrm{d}\theta \\ &\approx \sqrt{\pi} r_p \frac{k_p}{2} + \sqrt{\pi} r_p^2 \frac{k_p}{4\sqrt{2}R} = \sqrt{\pi} r_p \frac{k_p}{2}\left(1 + \frac{r_p}{2\sqrt{2}R}\right).\end{aligned} \tag{A12}$$

Numerical simulations consider a number of cells given by $N = \rho(\pi R^2 + n_p \Delta A)$.

#### b. Main text model parameters

To produce the figures in the main text (Fig. 6), we vary boundary perturbation parameters on the ranges $0 \leq h_p \leq 40\,\mu$m, $0 \leq k_p \leq 80\,\mu$m. We average the angular momentum over the last 100 h of the simulation. Since the model parameters are fixed, but we study orbiting stability for a varying ECM geometry, we extend the final simulation time to 500 h. After this time, all simulations appear to have converged to equilibrium. We choose adhesive parameters corresponding to the global orbiting regime $(f_a^{cc}, f_a^{lm}) = (0.2F_c, 0.2F_c)$. The rest of model parameters are taken as explained in Section 1 — see also Table S1.

### 4. Mosaic spheroids

#### a. Model description and parameters

Finally, we analyzed collective orbiting in a setting designed to mimic the mosaic spheroid experiments, in which simulations were initialized with a mixture of two cell types with distinct adhesion properties. Interestingly, our initial investigations showed that collective orbiting was relatively robust when the two cell types exhibited mismatched cell-cell adhesion but similar cell-matrix adhesion forces (Fig. S21). Observation of individual simulations suggested that this was likely because low adhesion ("Snail") cell are simply pushed by, and travel with, higher adhesion WT cells which otherwise move in a circular direction (Fig. S10). Guided by the experimental observation that Snail cells preferentially sort to the boundary and within the spheroid interior, we modified Snail cell model parameters such that those agents experience weaker cell-cell attraction and stronger cell-matrix adhesion forces than WT cells. Additionally, we imposed that Snail cells may isotropically sense the matrix. Interestingly, this mechanism proves to be important for disrupting orbiting (see SI for a discussion). Collective orbiting was robust to the introduction of relatively small fractions of Snail cells (up to ∼10%), with the angular order parameter roughly equal to $\langle L\rangle \sim 0.7 - 1.0$. Observation of individual simulations suggested that WT cells at the periphery maintain collective orbiting by simply migrating around relatively static Snail cells located at the periphery (Fig. 6GH). However, initializing simulations with larger Snail cell fractions caused the cell-matrix interface to become fully occupied by Snail cells, which impeded the ability of WT cells to sense and follow the boundary. This resulted in a sharp decrease in the angular order parameter to $\langle L\rangle < 0.2$ for Snail fractions above 15%. These simulations validate the importance of boundary curvature for coordinated orbiting within a parameter regime with intermediate cell-cell adhesion and weak cell-matrix adhesion. Further, an increasing fraction of obstructionist cells that occupy the boundary can also destabilize collective orbiting.

To describe the mosaic spheroid experiments, we consider a version of the mathematical model with two cell species. Both wildtype (WT) (1) and Snail cells (2) follow the same equations, but with different model parameters,

$$\frac{\mathrm{d}\mathbf{x}_i^{(1)}}{\mathrm{d}t} = \mathbf{v}_i^{(1)}\,,$$

$$\frac{\mathrm{d}\mathbf{v}_i^{(1)}}{\mathrm{d}t} = \left(\alpha - \beta\left|\mathbf{v}_i^{(1)}\right|^2\right)\mathbf{v}_i^{(1)} + \sum_{l\neq i,\, j=1}^{N_1} \mathbf{F}_{11}^{cc}(\mathbf{x}_l^{(1)} - \mathbf{x}_i^{(1)}) + \sum_{l=1}^{N_2} \mathbf{F}_{12}^{cc}(\mathbf{x}_l^{(2)} - \mathbf{x}_i^{(1)}) + \sum_{k=1}^{M} \mathbf{F}_1^{cm}(\mathbf{y}_k - \mathbf{x}_i^{(1)}, \mathbf{v}_i^{(1)})\,,$$

$$\frac{\mathrm{d}\mathbf{x}_j^{(2)}}{\mathrm{d}t} = \mathbf{v}_j^{(2)}\,,$$

$$\frac{\mathrm{d}\mathbf{v}_j^{(2)}}{\mathrm{d}t} = \left(\alpha - \beta\left|\mathbf{v}_j^{(2)}\right|^2\right)\mathbf{v}_j^{(2)} + \sum_{l=1}^{N_1} \mathbf{F}_{21}^{cc}(\mathbf{x}_l^{(1)} - \mathbf{x}_j^{(2)}) + \sum_{l\neq j,\, l=1}^{N_2} \mathbf{F}_{22}^{cc}(\mathbf{x}_l^{(2)} - \mathbf{x}_j^{(2)}) + \sum_{k=1}^{M} \mathbf{F}_2^{cm}(\mathbf{y}_k - \mathbf{x}_i^{(2)}, \mathbf{v}_i^{(2)})\,,$$

$$\frac{\mathrm{d}\mathbf{y}_k}{\mathrm{d}t} = \mathbf{0}\,,$$

where $N_1$ and $N_2$ are the number of WT and Snail cells respectively; $\mathbf{F}_{11}^{cc}$, $\mathbf{F}_{22}^{cc}$ represent adhesive-repulsive forces between the same type of cell; and $\mathbf{F}_{21}^{cc}$, $\mathbf{F}_{12}^{cc}$ represent cross-interaction forces.

TABLE S2. **Summary of two-species model and simulation parameters and their values used in numerical simulations.** The remaining model parameters (cell-cell repulsion, active-drag forces, and ECM geometry) are the same as in the one-species case (Table S1).

| Parameter | Values | Units | Meaning |
|---|---|---|---|
| $N_1$ | $\{85, 86, \ldots, 130\}$ | cells | number of WT cells |
| $N_2$ | $N - N_1$ | cells | number of Snail cells |
| $f_{a_{1,1}}^{cc}$ | $0.2F_c$ | $\mu$m/h$^2$ | cell-cell adhesion strength (WT-WT) |
| $f_{a_{1,2}}^{cc}$, $f_{a_{2,1}}^{cc}$ | 0 | $\mu$m/h$^2$ | cell-cell adhesion strength (WT-Snail) |
| $f_{a_{2,2}}^{cc}$ | 0 | $\mu$m/h$^2$ | cell-cell adhesion strength (Snail-Snail) |
| $f_{a_1}^{cm}$ | $0.2F_c$, $0.4F_c$ | $\mu$m/h$^2$ | cell-matrix adhesion strength (WT-ECM) |
| $f_{a_2}^{cm}$ | $0.8F_c$ | $\mu$m/h$^2$ | cell-matrix adhesion strength (Snail-ECM) |
| $\theta_{\max}^{(1)}$ | $\pi/3$ | rad | maximum cell-matrix alignment angle (WT) |
| $\theta_{\max}^{(2)}$ | $\pi$ (main text), $\pi/3$ (Fig. S21) | rad | maximum cell-matrix alignment angle (Snail) |

The forces, $\mathbf{F}_1^{cm}$ and $\mathbf{F}_2^{cm}$, model cell-matrix interactions for each cell type, similarly to the one-species case. These forces follow the expressions in Eqs. (A4)-(A6). We also note that active and drag forces are identical for both cell types. We summarize the chosen model parameters to produce Fig. 6 in the main text in Table S2. As in the previous section, we solve numerically the model until a final time of 500 h, and record angular momentum using the last 100 h of the simulation. Cells are initialized as in the one-species case.

### 5. Additional simulations

In this section, we present additional numerical simulations of the model, where we test the impact of varying different model parameters.

#### a. *Cell-matrix alignment angle ($\theta_{\max}$)*

We begin by exploring the impact of the cell-matrix alignment angle $\theta_{\max}$(see Fig. S18), which sets the maximum angle between the cell velocity and the relative position to the spheroid boundary (Eq. (A5)). As expected intuitively, we find that for small values of $\theta_{\max}$, orbiting is disrupted easily. The angular momentum

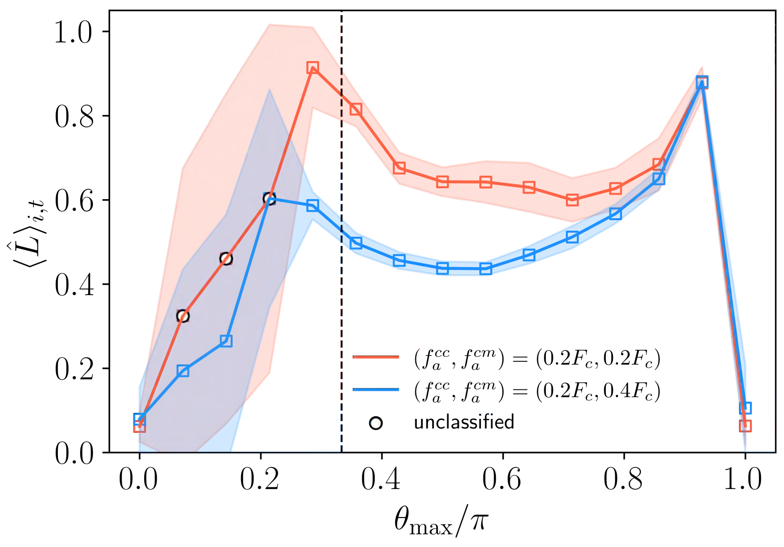


FIG. S18. **Impact of cell-matrix alignment on collective orbiting.** All parameters are fixed except for $\theta_{\max}$ in Eq. (A5), which is varied. Parameter values are taken from Section 1, with cell-matrix and cell-cell adhesion parameters chosen to represent the global (red) and peripheral orbiting (blue) regimes. Angular momentum is recorded over $t \in [200, 300]$ h and averaged across thirty simulations. Shaded regions indicate one standard deviation. The black dashed line denotes $\theta_{\max} = \pi/3$, which is the value used in the main text.

parameter peaks at a value close to 1 (for comparable cell-cell and cell-matrix adhesion parameters), and for values close to those used in the main text ($\theta_{\max} = \pi/3$). Numerical results remain consistent with the main text for larger values of the maximum cell-matrix alignment angle ($\theta_{\max} \sim 0.3\pi - 0.8\pi$). Interestingly, when $\theta_{\max}$ is close to $\pi$, meaning that cells can adhere to the matrix from nearly any direction except directly behind them, global orbiting appears to be more stable. This may be due to the increased range over which cells can detect and interact with the matrix, enhancing their ability to align with the boundary. However, in the limit of isotropic interactions, orbiting is completely disrupted ($\langle \hat{L} \rangle_{i,t} \sim 0$), highlighting the crucial role of anisotropy in cell-matrix interactions for the emergence of orbiting.

#### *b. Active-drag forces ($\alpha$, $\beta$)*

The main text uses a self-propulsion parameter of $\alpha = 0.1$ h$^{-1}$, setting a timescale for speed readjustments due to active-drag forces at approximately 10 hours. This choice yields realistic cell speeds. To explore the effect of this parameter on orbiting behavior (Fig. S19), we vary $\alpha$ by an order of magnitude ($\alpha = 10^{-2}$, $10^0$ h$^{-1}$) while adjusting the friction coefficient, $\beta$, to keep the equilibrium speed in the absence of cell-matrix interactions, $\sqrt{\alpha/\beta} = u$, constant.

In both cases, the phase diagram reveals regions of global orbiting, peripheral orbiting, and arrested motion (Fig. S19). The timescales to reach these states remain largely unchanged, supporting our numerical findings that they are primarily governed by the strength of the interaction kernel. Additionally, Fig. S19 suggests that cell speeds decrease with increasing self-propulsion strength, consistent with equilibrium speed being determined by the balance between active-drag and cell-matrix forces.

While all three cases exhibit similar behavior, when $\alpha = 10^{-2}$ h$^{-1}$, a larger portion of the phase diagram corresponds to higher values of the angular momentum order parameter, indicating a global orbiting state. This may result from cell-matrix interactions dominating over active-drag forces, allowing cells to synchronize within the spheroid. Conversely, stronger self-propulsion accelerates the onset of arrest in the strong cell-cell adhesion regime.

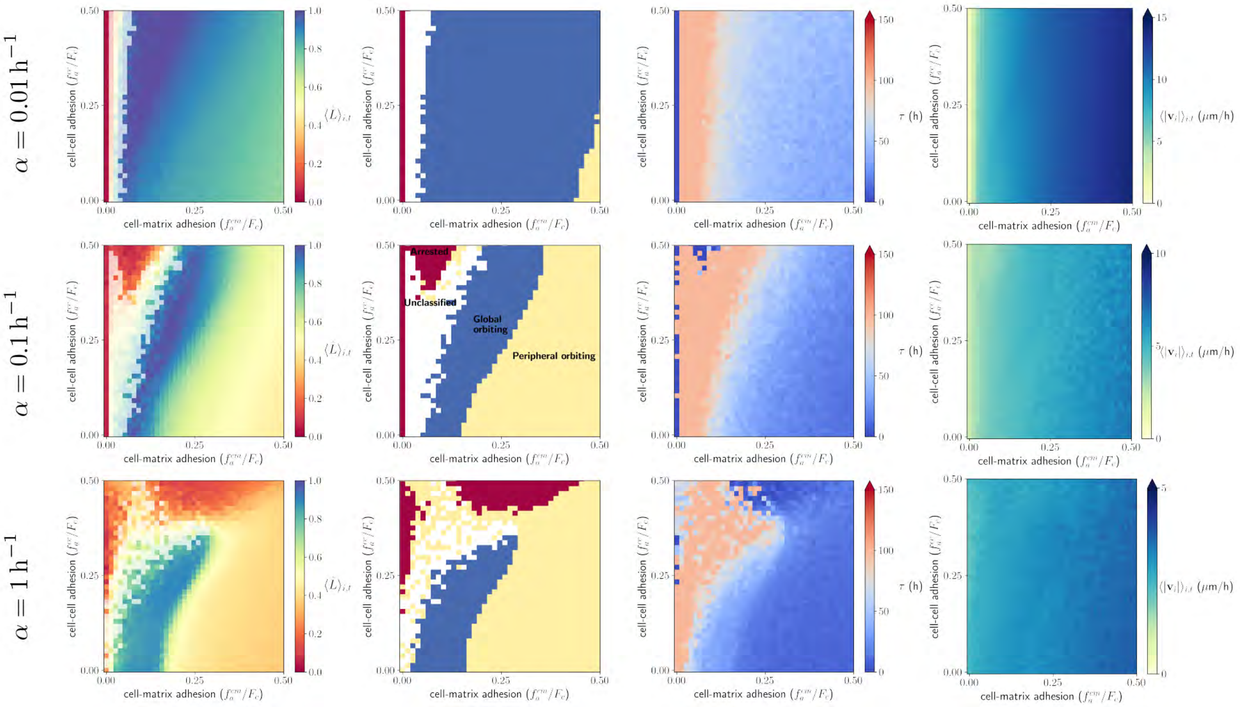


FIG. S19. **Impact of active-drag forces on collective orbiting.** All parameters are fixed except for $\alpha$ (and $\beta = \alpha/u^2$ to keep $u$ constant), which is varied. Parameter values are taken from Section 1. Angular momentum is recorded over $t \in [200, 300]$ h and averaged across fifty simulations. The columns, from left to right, represent the normalized angular momentum phase diagram, the discretized phase diagram of orbiting regimes, the timescale to orbiting ($\tau$, Eq. (A9)), and the mean cell velocity.

#### c. *Three boundary perturbations ($n_p = 3$)*

We further examined the effect of three outward-oriented Gaussian perturbations of the ECM boundary and found a similar destabilizing effect on collective orbiting as in the case of two perturbations. Beyond a critical perturbation size—approximately one cell diameter in height and two cell diameters in width—orbiting was disrupted (Fig. S20). The phase diagram closely resembles the two-perturbation case, reinforcing that multiple sharp boundary features make coordinated migration more difficult by reducing the length of continuously curved boundary available for collective motion.

#### d. *Differential adhesion and orbiting in two-species spheroids ($\theta^{(2)}_{\max}$)*

We also investigated the case where Snail cells exhibit directionality when sensing the matrix, implemented by setting the same maximum cell-matrix alignment angle as WT cells ($\theta^{(2)}_{\max} = \theta^{(1)}_{\max} = \pi/3$). In this scenario, orbiting remains stable in mixed cell populations, as both cell types are capable of orbiting independently (Fig. S21). Interestingly, these simulations reveal classical sorting patterns driven by differential adhesion [14], considering both cell-cell and cell-matrix interactions. Since Snail cells exhibit weaker cell-cell adhesion but stronger cell-matrix adhesion, they preferentially localize at the boundary while maintaining peripheral

---

[14] Armstrong, N.J. et al. J. Theor Biol. 243(1), 98-113 (2006); Buttenschön, A. et al. Bull Math Biol. 86(11), 129 (2024); Carrillo, J.A.., et al. J. Theor Biol. 474, 14-24 (2019); Falcó, C. et al. *SIAM J. Appl. Math.* 84(3), S17-42 (2023).

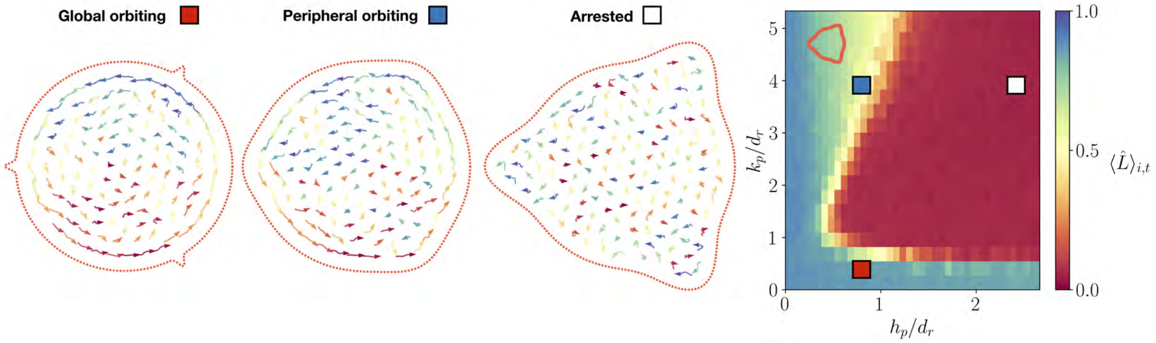


FIG. S20. **Orbiting is disrupted by three evenly spaced boundary perturbations.** Representative snapshots of global orbiting (red square), peripheral orbiting (blue square) and arrested migration (white square) with three boundary perturbations, and the corresponding phase diagram depicting how the angular order parameter phase diagram responds to different perturbation heights and widths.

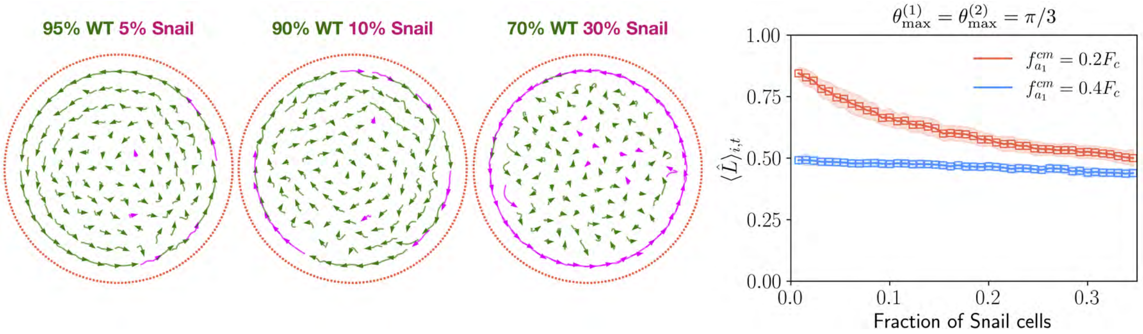


FIG. S21. **Orbiting and differential adhesion in two-species spheroids.** Representative snapshots of numerical simulations with increasing percentages of Snail populations are shown. The graph on the right depicts how the average normalized angular momentum over the last 100 hours of the simulation varies with the fraction of snail cells. Red squares denote simulations in which the cell-matrix adhesion parameter for WT cells, $f^{cm}_{a_1}$, is equal to $0.2F_c$, while blue squares indicate simulations in which the same parameter is set equal to $0.4F_c$ (see Table S2 for more details). Shaded regions indicate one standard deviation.

orbiting. These findings highlight that disrupting orbiting requires a change in the directionality of cell-matrix interactions, as demonstrated in the main text.
